# Low-dose Perampanel rescues cortical gamma dysregulation associated with parvalbumin interneuron GluA2 upregulation in epileptic *Syngap1*^+/-^ mice

**DOI:** 10.1101/718965

**Authors:** Brennan J. Sullivan, Simon Ammanuel, Pavel A. Kipnis, Yoichi Araki, Richard L. Huganir, Shilpa D. Kadam

**Affiliations:** Neuroscience Laboratory, Hugo Moser Research Institute at Kennedy Krieger, Baltimore, MD 21205, USA; School of Medicine, University of California, San Francisco, 505 Parnassus Avenue, San Francisco, CA, 94143, USA; Department of Neuroscience, Kavli Neuroscience Discovery Institute, Johns Hopkins University School of Medicine, Baltimore, MD 21205, USA; Department of Neurology, Johns Hopkins University School of Medicine, Baltimore, MD 21205, USA

**Keywords:** Myoclonic seizures, sleep cycles, NREM, REM, interictal spikes, gamma oscillations, parvalbumin interneurons, AMPA receptors, Perampanel

## Abstract

Loss-of-function *SYNGAP1* mutations cause a neurodevelopmental disorder characterized by intellectual disability and epilepsy. SYNGAP1 is a Ras-GTPase-activating protein that underlies the formation and experience-dependent regulation of postsynaptic densities. The mechanisms that contribute to this proposed monogenic cause of intellectual disability and epilepsy remain unresolved. Here, we establish the phenotype of the epileptogenesis in a *Syngap1^+/-^* mouse model using 24h video electroencephalogram/electromyogram (vEEG/EMG) recordings at advancing ages. A progressive worsening of clinically-similar seizure phenotypes, interictal spike frequency, sleep dysfunction, and hyperactivity was identified in *Syngap1^+/-^* mice. Interictal spikes emerged predominantly during NREM in 24h vEEG of *Syngap1^+/-^* mice. Myoclonic seizures occurred at behavioral-state transitions both in *Syngap1^+/-^* mice and during an overnight EEG from a child with *SYNGAP1* haploinsufficiency. In *Syngap1^+/-^* mice, EEG spectral power analyses identified a significant loss of cortical gamma homeostasis during behavioral-state transitions from NREM to Wake and NREM to REM. The loss of gamma homeostasis was associated with a region- and location-specific significant increase of GluA2 AMPA receptor subunit expression in the somas of parvalbumin-positive (PV+) interneurons. Acute dosing with Perampanel, an FDA approved AMPA antagonist significantly rescued cortical gamma homeostasis, identifying a novel mechanism implicating Ca^2+^ impermeable AMPARs on PV+ interneurons underlying circuit dysfunction in *SYNGAP1* haploinsufficiency.

## Introduction

*SYNGAP1* codes for synaptic Ras GTPase activating protein 1 (SYNGAP1), a Ras-GAP critical for the formation of postsynaptic densities (PSDs) and the experience-dependent AMPA receptor (AMPAR) insertion that underlies synaptic plasticity^1–5^. Mutations in *SYNGAP1* are prevalent in patients with schizophrenia, intellectual disability (ID), and autism spectrum disorder^6–8^. Loss-of-function *SYNGAP1* mutations result in haploinsufficiency and cause mental retardation type 5 (MRD5, OMIM#612621), a severe distinct generalized developmental and epileptic encephalopathy with ID, ataxia, severe behavioral problems, and a risk for autism^6–12^.

The majority of patients with MRD5 have refractory epilepsy^12^. Poor quality sleep is highly prevalent in patients with neurodevelopmental disorders (NDDs) and epilepsy^13, 14^. The relationship between epilepsy and dysfunctional sleep is a major focus of ongoing clinical and pre-clinical research^13–15^. For example, Rett syndrome (RTT) is an NDD with comorbidity of epilepsy and sleep dysfunction^16, 17^. Research in pre-clinical models of RTT, as well as patients with RTT has identified translatable quantitative EEG (qEEG) biomarkers^17, 18^. There is an urgent need to develop robust biomarkers for NDDs, as bench-to-bedside therapeutic strategies are severely hindered without validated quantitative outcome measures.

Patients with *SYNGAP1* mutations predominantly present with myoclonic, absence, or tonic-clonic seizures^8, 12^. Clinical cohort studies reveal progressive epilepsy followed by spontaneous remission of epilepsy in a subset of patients with *SYNGAP1* mutations in their late teens^12^. Children with epileptic encephalopathies have a slow developmental regression that is primarily due to seizures, interictal spikes (IISs), or cortical dysrhythmia identified on EEG^19, 20^. Case reports indicate that adult patients demonstrate gradual decline in cognitive abilities^21^. Perampanel (PMP), a recently FDA approved AMPAR antagonist has shown significant promise for multiple seizure types in idiopathic generalized epilepsies including absence^22^. Its use in infants with epilepsy is in clinical trial^23^. However, the role of AMPAR antagonists such as PMP in SYNGAP1 haploinsufficiency related seizures and ID is unknown.

Mice with Syngap1 haploinsufficiency (*Syngap1^+/-^*) are relevant translational models presenting with learning and memory deficits, abnormal dendritic spine dynamics, cortical hyperexcitability, and precocious unsilencing of thalamocortical synapses during development^24–26^. Currently, no studies have investigated the epileptogenesis, sleep, and underlying mechanisms associated with MRD5. An understanding of the mechanisms that promote spontaneous seizures and network dysfunction can help accelerate the discovery of novel therapeutic strategies. For this purpose, a *Syngap1^+/-^* loss-of-function mouse model^27^ (exon 7 and 8 deletions) underwent 24h video-vEEG/EMG recordings at advancing ages (P60-P120). A subsequent 24h EEG investigated the effect of low-dose PMP on EEG identified biomarkers.

## Results

### *Syngap1^+/-^* mice have recurrent spontaneous seizures

Patients with pathogenic *SYNGAP1* mutations have epilepsy^12^. The predominant seizure phenotypes are absence, eyelid myoclonia with absences, and myoclonic seizures. The myoclonic seizures are distinguished from atypical absences by pronounced myoclonic jerks and a global spike wave discharge ∼3Hz on EEG^12^. Continuous 24h vEEG/EMG recordings at temporally advancing ages were utilized to investigate and identify spontaneous seizures in 50% (n/n; 4/8) of *Syngap1^+/-^* mice (Fig. 1). At P60 all seizures were myoclonic (5/5); 80% (4/5) of these seizures started in non-rapid eye movement (NREM) at NREM-Wake transitions (Fig. 1A). Myoclonic seizures were 34.4±5.9s in duration, with rhythmic spike-wave discharges occurring ∼3-4Hz frequency (Fig. 1A). The myoclonic seizures were distinguished by their time-locked myoclonic jerks recorded on EMG with video confirmation of head, neck, and upper shoulder myoclonic jerks (see Supplemental Video 1).

**Fig. 1.**
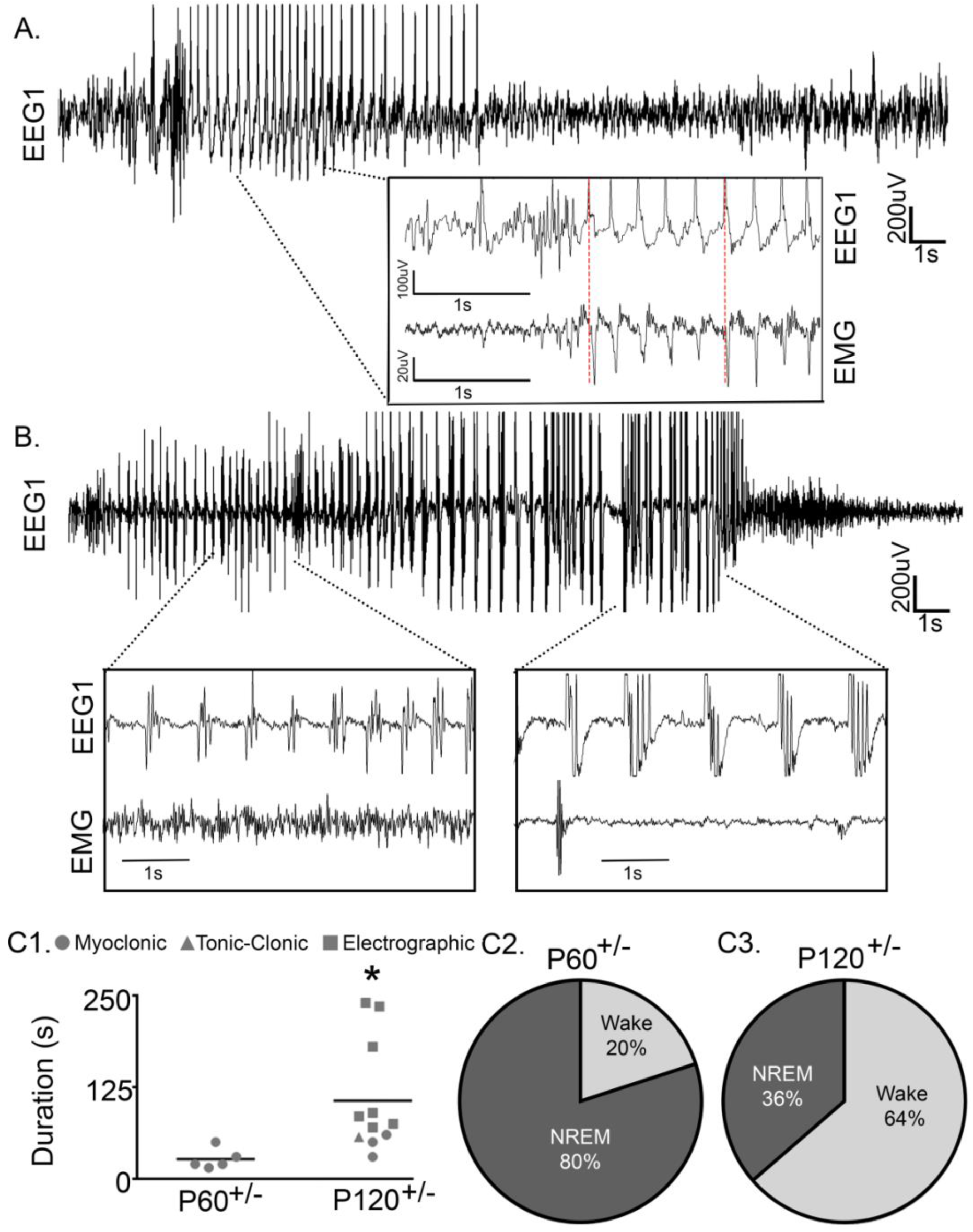
*Syngap1*^+/-^ mice had recurrent spontaneous seizures. **(A)** A representative EEG trace (0.5-50Hz) of a spontaneous myoclonic seizure at P60 during NREM sleep. Prototypical time-locked myoclonic jerks were recorded on EMG (see Supplementary Video 1). **(B)** A representative spontaneous tonic-clonic seizure at P120 during NREM sleep. **(C1)** The average duration of seizures significantly increased from P60 to P120 (*t-test*, t_14_=2.307, p=0.037). **(C2)** At P60 all seizures were myoclonic and the majority occurred during NREM (4/5). **(C3)** At P120, all myoclonic (3/11) and tonic-clonic (1/11) seizures occurred during NREM, whereas all electrographic seizures occurred during wake (7/11) (for the electrographic seizure trace see Supplementary Figure 2).

Multiple seizure phenotypes were identified in P120^+/-^ mice consistent with the latest clinical reports^12, 28^. Myoclonic, generalized tonic-clonic, and electrographic seizures were observed (Fig. 1B, 1C1, and Supplemental Fig. 2). The majority of seizures were electrographic seizures emerging during wake (n/n; 7/11), while all myoclonic seizures occurred during transitions from NREM to Wake (3/11) a common feature for several epilepsy syndromes such as juvenile myoclonic epilepsy^29^. A single tonic-clonic seizure occurred during NREM (1/11). Moreover, all myoclonic and tonic-clonic seizures occurred during NREM in P120^+/-^ mice, whereas all electrographic seizures occurred during wake (Fig 1C3). In P60^+/-^ mice, all seizures were myoclonic and 4/5 occurred during NREM (Fig. 1C2). From P60 to P120, the average seizure duration significantly increased from 27±0.6 to 106±22.7s respectively (Fig. 1C1). Together, these data suggest that myoclonic seizures had a propensity to emerge during NREM and were of significantly shorter duration than all other seizure phenotypes.

### Recurrent generalized seizures in human *SYNGAP1* haploinsufficiency show affinity for transition-states and clustering in NREM

To evaluate the translational value of qEEG analysis of 24h EEGs in *Syngap1^+/-^* mice, we initiated an IRB-approved study to acquire copies of 24h EEG recordings from children diagnosed with pathogenic *SYNGAP1* mutation related epilepsy^30^. A 20h continuous EEG which captured an entire cycle of overnight sleep in a 3 year old boy with a *de novo* intronic variant in *SYNGAP1* associated with myoclonic and atonic absence seizures (Fig. 2) was analyzed. The global seizures were bursts of 3-4Hz spike and slow waves that lasted ∼0.5-3s (Fig. 2A). The EEG was sleep-scored for wake, NREM, and REM using video and qEEG spectral power analysis similar to our previously published protocols^17^. Ictal events occurred both in wake and sleep. All ictal events observed during sleep occurred in NREM and specifically at Wake-NREM and NREM-REM transitions (Fig. 2B). We identified a proclivity for clustering of ictal events (i.e. inter-ictal durations <5min) that was predominant during NREM (Fig. 2C & D). These analyses for the first time identify clustering of ictal-events during NREM in a child with *SYNGAP1* related ID. The propensity of ictal events to occur at transition-states is a novel finding and may be a biomarker for underlying mechanisms.

**Fig. 2.**
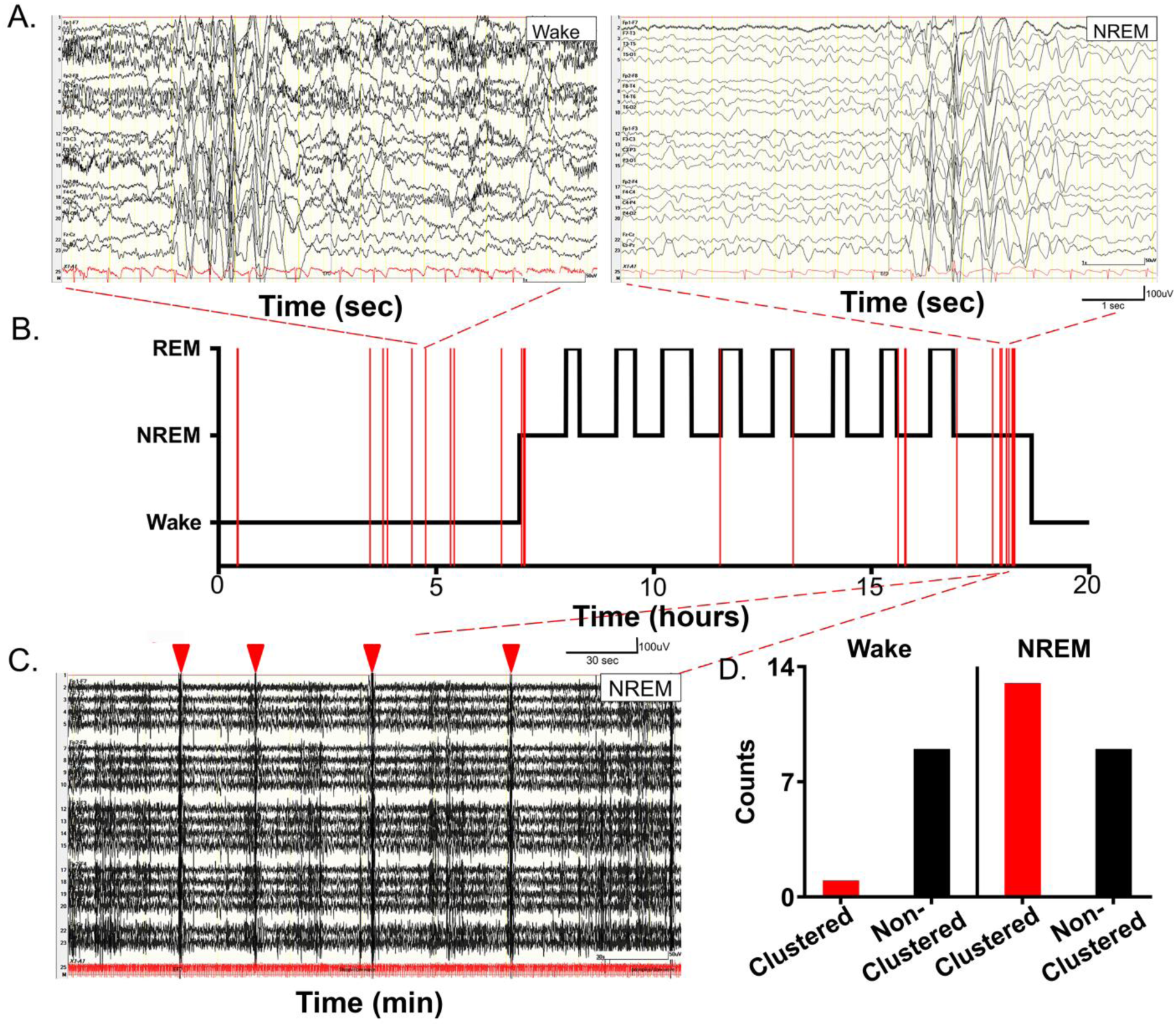
Ictal events over a 20h EEG recording period spanning wake and sleep in a 3 year old boy with pathogenic *SYNGAP1* mutation. (X/350 words) (A) Identical 3Hz short duration epileptiform events during wake and NREM sleep. **(B)** Hypnogram for the 20h period plotted using video and delta power during NREM and REM, determined by fast-Fourier transformation after conversion to EDF format, as previously reported (Ammanuel et. al; 2015)). All ictal events plotted as a raster plot overlying the hypnogram. Obvious clustering of events identified during NREM sleep. **(C)** 5min EEG trace showed four clustered events (red arrowheads) occurring within a 5 min period during the last NREM cycle before wake. Note ictal events occurring at transition points between wake and sleep and REM-NREM. **(D)** Bar graphs grouping clustered vs. non-clustered events for wake and NREM showed proclivity of ictal clustering during NREM. Seizure rates normalized per hour of EEG recording for the raster were wake=1.27/h; NREM=2.68/h and REM=0/h for this recording.

### Progressive alterations in macro-sleep architecture

Sleep dysfunction is a common feature of NDDs and is also reported in patients with *SYNGAP1* mutations^12, 21^. Quantitative assessments of the sleep problems in patients with *SYNGAP1* mutations are lacking; therefore, a robust assessment is warranted^30^. In this study, the WT^+/+^ sleep architecture was ultradian with 15.38±2.2 sleep cycles occurring over 24h (Fig. 3A). WT^+/+^ mice spent ∼60% of the 24h period awake. During light vs. dark phase, WT^+/+^ mice spent ∼50% and ∼80% of the time awake (Fig. 3A - B2), respectively, demonstrating the expected nocturnal predominance of activity in WT^+/+^ mice.

**Fig. 3.**
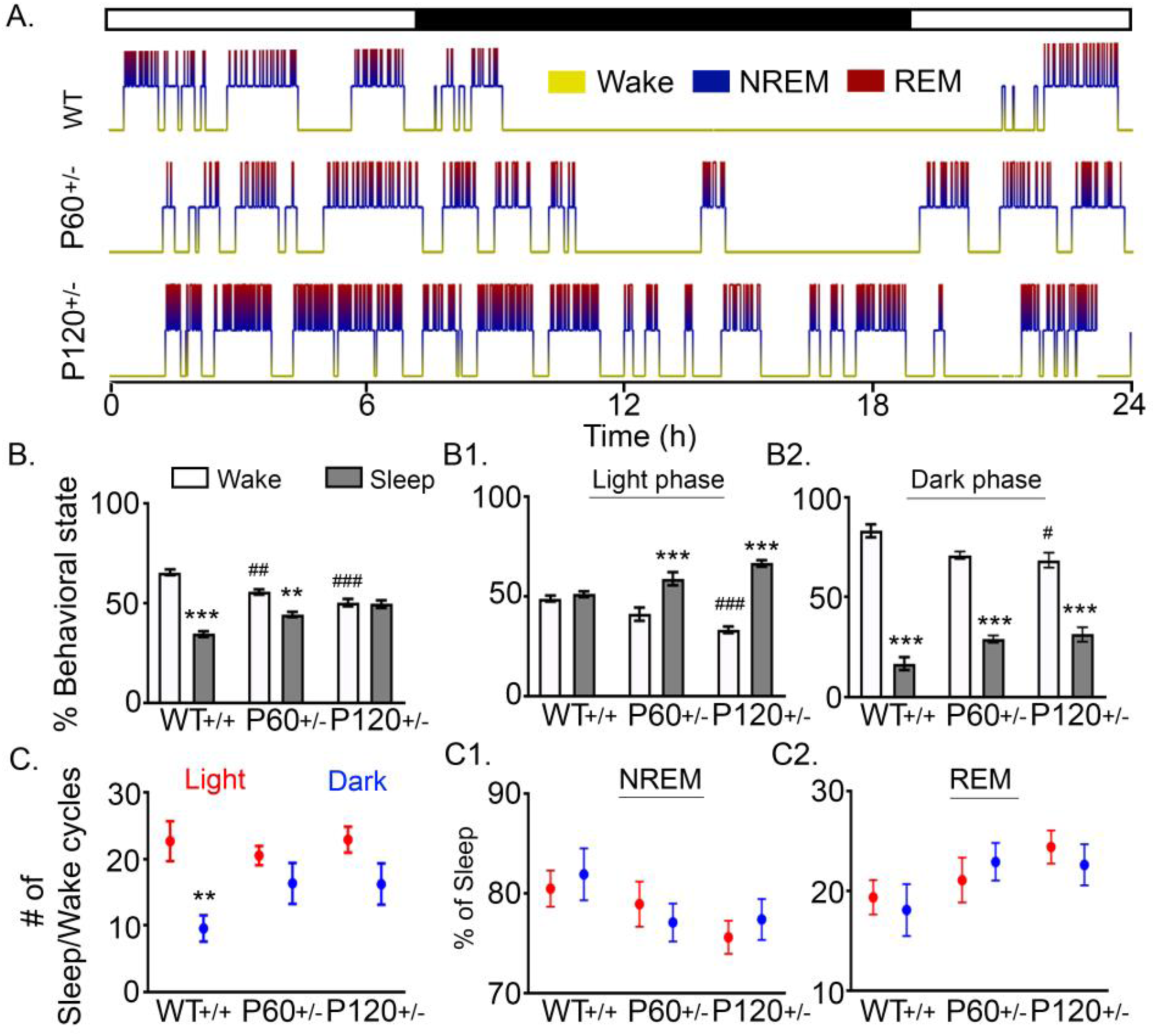
Progressive alterations in macro-sleep architecture. **(A)** 24h hypnograms of nocturnal ultra-radian rhythms in WT^+/+^, P60^+/-^, and P120^+/-^ mice. Regardless of age, all WT^+/+^ mice had consistent macro and micro-sleep architectures (Supplemental Fig. 3). **(B)** All WT^+/+^ mice spent significantly less time sleeping than awake (WT^+/+^ Wake vs. WT^+/+^ Sleep: 2-way ANOVA, F_2,42_=42.03, P<0.0001, p<0.0001). P60^+/-^ mice also spent significantly less time sleeping than awake (P60^+/-^ Wake vs. P60^+/-^ Sleep: 2-way ANOVA, F_2,42_=42.03, P<0.0001, p=0.002), however P120^+/-^ mice lacked any significant differences between wake and sleep durations. Between groups, time spent awake significantly decreased in P60^+/-^ (WT^+/+^ Wake vs. P60^+/-^ Wake: 2-way ANOVA, F_2,42_=42.03, P <0.0001, p=0.003) and P120^+/-^ mice (WT^+/+^ Wake vs. P120^+/-^ Wake: 2-way ANOVA, F_2,42_=42.03, P <0.0001, p<0.0001). **(B1)** During the 12h light phase both P60^+/-^ and P120^+/-^ mice spent more time asleep than awake (P60^+/-^ Light Wake vs. P60^+/-^ Light Sleep: 2-way ANOVA, F_2,42_=29.02, P<0.0001, p<0.0001; P120^+/-^ Light Wake vs. P120^+/-^ Light Sleep, p<0.0001). During the light phase, the amount of time P120^+/-^ mice spent awake was significantly less than WT^+/+^ (WT^+/+^ Light Wake vs. P120^+/-^ Light Wake: F_2,42_=29.02, P <0.0001, p<0.0001). **(B2)** WT^+/+^, P60^+/-^, and P120^+/-^ mice all spent significantly less time asleep than awake during the dark phase (WT^+/+^ Dark Wake vs. WT^+/+^ Dark Sleep: 2-way ANOVA, F_2,42_=10.95, P=0.0001, p<0.0001; P60^+/-^ Dark Wake vs. P60^+/-^ Dark Sleep: p<0.0001, P120^+/-^ Dark Wake vs. P120^+/-^ Dark Sleep: p<0.0001). Between groups, P120^+/-^ mice spent significantly less time awake during the dark phase than WT^+/+^ (WT^+/+^ Dark Wake vs. P120^+/-^ Dark Wake: p<0.0001, P120^+/-^ Dark Wake vs. P120^+/-^ Dark Sleep: p=0.0256). **(C)** Only WT^+/+^ mice had significantly fewer cycles (transitions between sleep and wake) during the dark phase than the light phase (WT^+/+^ Light cycles vs. WT^+/+^ dark cycles: 2-way ANOVA, F_2,42_=1.547, P=0.0001, p=0.0066. **(C1 & C2)** Although this significant diurnal difference in the number of sleep/wake cycles detected in WT^+/+^ did not occur t in P60^+/-^ and P120^+/-^ mice, the micro-architecture of both NREM and REM cycles during sleep was not significantly different. (p<0.05 *, p<0.01 **; P120 p<0.001 ***; p<0.05 #, p<0.01 ##; p<0.001 ###, post-hoc Bonferroni).

Similar to WT^+/+^ mice, P60^+/-^ mice also spent significantly more time awake than asleep. In contrast, the P120^+/-^ mice lacked any significant differences between time spent awake and asleep (Fig. 3B). Between groups, P60^+/-^ and P120^+/-^ mice spent less time awake than WT^+/+^ mice over the 24h recording. This decrease in wake duration (and subsequent increase in sleep duration), was apparent during the light phase (Fig. 3B1). WT^+/+^ mice did not demonstrate any differences between time spent awake and asleep during the light phase. In contrast, P60^+/-^ and P120^+/-^ mice spent significantly less time awake than asleep (Fig. 3B1). Compared to WT^+/+^ mice, P120^+/-^ mice spent significantly less time awake.

During the dark phase, WT^+/+^ mice spent significantly more time awake than asleep. Nocturnal activity in WT^+/+^ mice included nest building, exploration, and grooming behaviors. All mice regardless of genotype or age spent a significantly greater amount of time awake than asleep during the dark phase (Fig. 3B2). However, compared to WT^+/+^ mice, P120^+/-^ mice spent significantly more time asleep in the dark phase (Fig. 3B2). These data suggest that P120*^+/-^* mice spent significantly more time asleep than WT^+/+^ mice.

In WT^+/+^ mice, there were significantly fewer sleep/wake cycles (transitions from wake to sleep) during the dark phase (9.6±2.0 cycles) compared to the light phase (22.67±9.6 cycles, Fig. 3C). During the dark phase, both P60^+/-^ and P120^+/-^ mice had more sleep/wake cycles (16.33±3.1 and 16.22±3.1 cycles, respectively) than WT^+/+^ mice (9.6±2.0 cycles), however this increase was not significant. Sleep/wake cycles during the light phase were not significantly different (Fig. 3C). In P60^+/-^ and P120^+/-^ mice, the difference between the numbers of sleep/wake cycles during light vs. dark phase was lost due to these alterations (Fig. 3C).

WT^+/+^ mice spent 6.99±0.3h in NREM and 1.76±0.2h in REM during the 24h recording. Progressive alterations to macro-sleep architecture in P60^+/-^ and P120^+/-^ mice led to an increase in the total duration of NREM (8.6±0.3 and 9.3±0.2h, respectively) and REM (2.4±0.3 and 3.00±0.3h, respectively) when compared to WT^+/+^ mice. The microarchitecture of sleep (the percentages of NREM and REM that constitute sleep) in WT^+/+^ mice was 80.3±1.7% NREM and 19.7±1.9% REM during the 24h recording (Fig. 3C1 & C2). P60^+/-^ and P120^+/-^ slept more than WT^+/+^ mice, yet their microarchitecture of sleep was not significantly different compared to WT^+/+^ mice (Fig. 3C1 & C2). Therefore, all groups maintained a sleep microarchitecture of ∼80% NREM and ∼ 20% REM.

### IISs predominantly occurred in NREM sleep

Uncontrolled epilepsy is at least partially responsible for cognitive regression in children with epileptic encephalopathies^19, 20^. IISs are reliable biomarkers of epileptogenesis that frequently occur on the EEGs of patients with epilepsy^31^. *Syngap1^+/-^* mice (n/n=8/8) presented IISs during both sleep when delta power and synchrony are high, and during wake when delta power and synchrony are relatively low (Fig. 4A).IISs significantly increased in frequency at P120*^+/-^* when compared to WT^+/+^ mice (1-way ANOVA, F_2,19_=5.27, P=0.015, p=0.013). This significant increase was driven by IISs that occurred during sleep (Fig. 4B) and was driven by light phase sleep (Fig. 4C). However, IISs during both light and dark phase sleep progressively increased in P120^+/-^ mice (Fig. 4C). The frequency of IISs per hour of NREM and REM sleep was significantly greater in P120^+/-^ mice (Fig. 4D). Therefore, taken together these data demonstrate that the overall number of IISs during the 24h recording period increased in P120^+/-^ mice and these IISs mostly occurred during sleep in the light phase. Furthermore, even though the total duration of sleep increased significantly in P120^+/-^ mice, (Fig. 3B) the rate of IIS occurrence per hour of NREM and REM sleep was significantly greater in P120^+/-^ mice compared to WT^+/+^ (Fig. 4D).

**Fig. 4.**
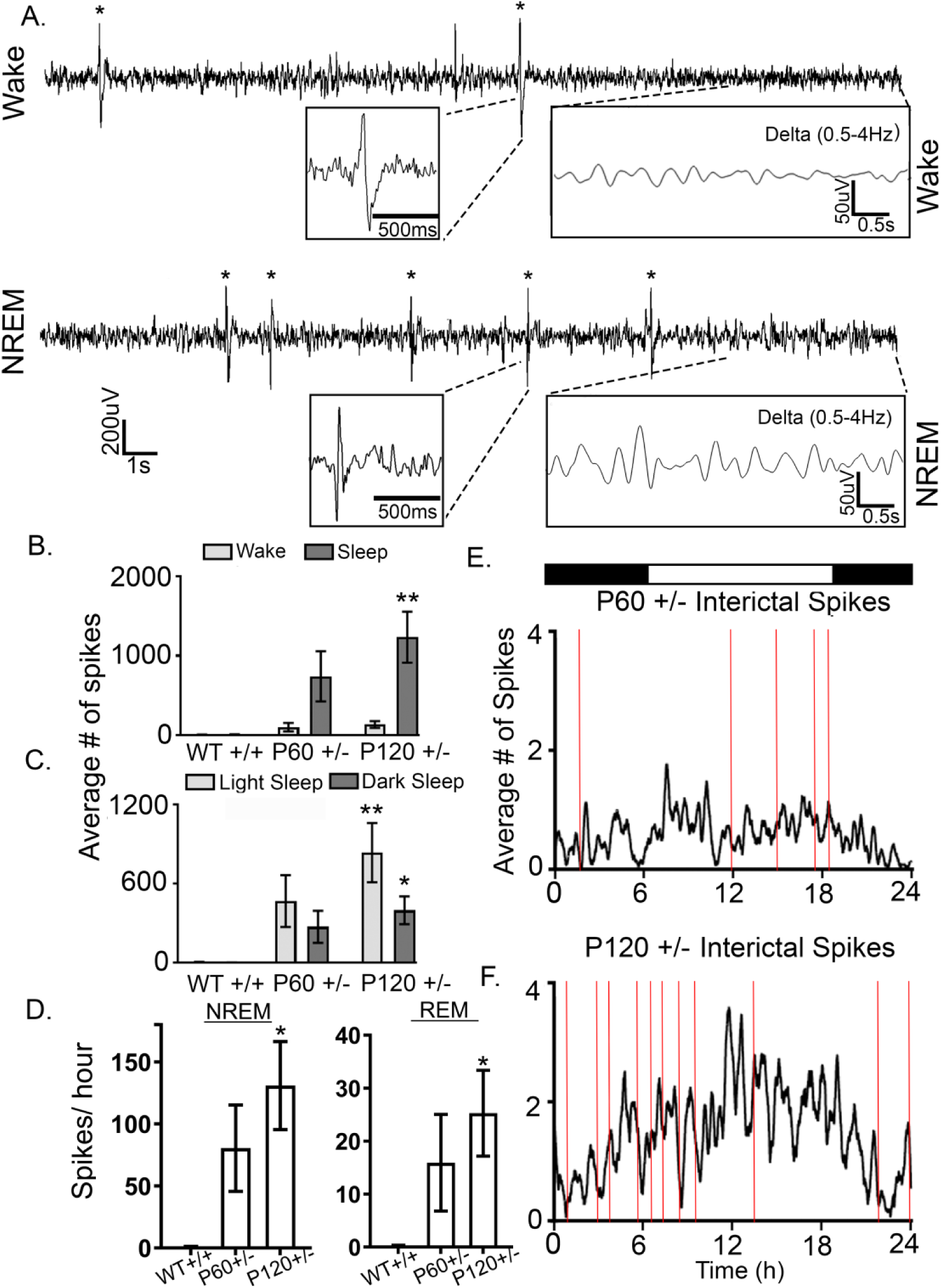
Progressive interictal spiking during NREM. **(A)** 30s EEG trace (0.5-50Hz) during wake of interictal spikes (n=2) with low amplitude delta power (0.5Hz-4Hz). During NREM sleep, a 30s EEG trace demonstrated interictal spikes (n=5) with high amplitude delta power (0.5Hz-4Hz). **(B)** Number of spikes during sleep significantly increased in P120^+/-^ compared to WT mice (WT^+/+^ Sleep Spikes vs. P120^+/-^ Sleep Spikes: 1-way ANOVA, F_2, 19_=5.36, P=0.014, p=0.012) **(C)** Number of spikes significantly increased during both light sleep and dark sleep in P120^+/-^ mice (WT^+/+^ vs. P120^+/^: 1-way ANOVA, Light Sleep Spikes: F_2,19_=5.386, P=0.014, p=0.012, Dark Sleep Spikes:F_2,19_=4.838, P=0.02, p=0.018). **(D)** Interictal spikes significantly increased with age during NREM (WT^+/+^ NREM spikes vs. P120^+/-^ NREM spikes: 1-way ANOVA, F_2, 19_=5.26, P=0.015, p=0.013) and REM (WT^+/+^ vs. P120^+/^:1-way ANOVA, F_2, 19_=3.458, P=0.052, p=0.050). **(E & F)** Interictal spikes (black traces) and seizure frequency (red lines) over 24h. (p<0.05 *, p<0.01 **; p<0.001 ***; post-hoc Bonferroni).

Emerging research utilizing intracranial recordings from epilepsy patients has demonstrated that regulation of brain activity occurs on long timescales^32^. Daily patterns of seizure occurrence demonstrate circadian rhythmicity and clustering organization^32^. In this study, we report the relationship between seizure occurrence and frequency distribution of IISs over 24h for all mice (Fig. 4E & 4F). Grouped data of IIS frequency averaged over 24h showed an escalation in IIS frequency at the transition between the dark and light phases associated with increased incidence of seizures in P120^+/-^ mice which was not apparent in P60^+/-^ mice. Antagonizing AMPARs acutely with Perampanel (PMP; 2mg/kg I.P. at 10AM and 6PM) prevented seizures during the 24h EEG. However, PMP had a modest effect on IIS frequency (Supplementary Fig. 4). In summary, the data suggest that IIS and seizure occurrence was greater during sleep in *Syngap1^+/-^* mice.

### Hyperactivity worsened with age

Video tracking software allows for high specificity and sensitivity when tracking mouse motor activity over the 24h recordings. Total distance traveled by each mouse over a24h period (Fig. 5A1 - A3) was used to identify hyperactivity in *Syngap1^+/-^* mice. Compared to WT^+/+^ mice (267.92±46.8m over 24h), P120^+/-^ mice traveled greater distances (479.75±46.1m over 24h) that was not detected in the P60^+/-^ mice (288.22±32m over 24h; Fig. 5A2 & A3). Since P120^+/-^ mice spent more time asleep than WT^+/+^ mice (Fig. 3), the distance traveled by each mouse was normalized to the duration of time spent asleep. The running average of all mice in each group highlighted increased motor activity of P120^+/-^ mice at the end of the dark phase when the majority of nesting behavior occurs^33^ (Fig. 5B). The normalized distance traveled by P120^+/-^ mice was significantly greater than WT^+/+^ and P60^+/-^ mice (Fig. 5C). Taken together these data suggest that P120^+/-^ mice spent more time asleep but were hyperactive when awake, as they travelled twice the distance compared to WT^+/+^ mice over 24h. These results are consistent with recent reports of home cage hyperactivity in male *Syngap1^+/-^* mice^34, 35^.

**Fig. 5.**
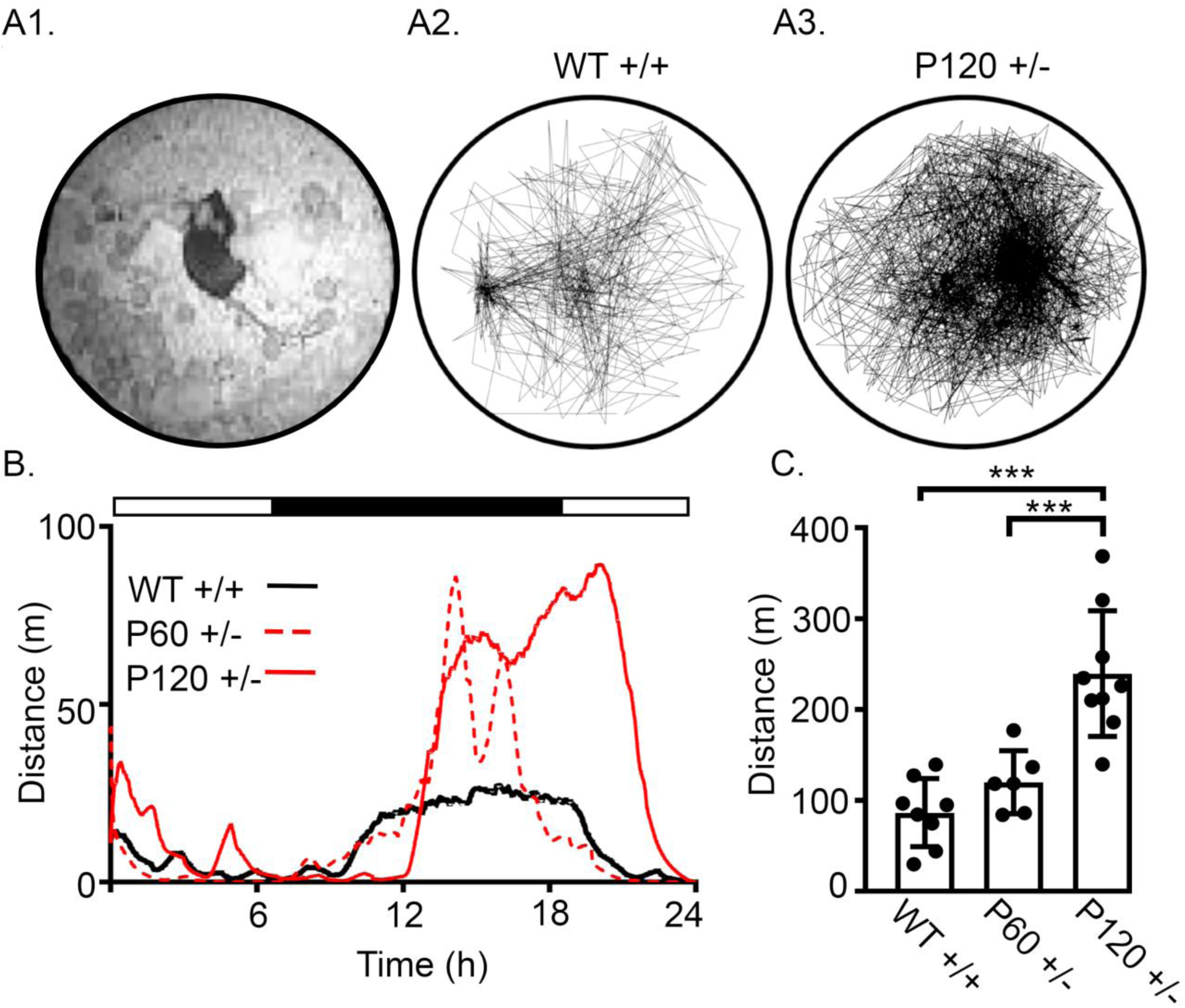
Progressive increase in activity during 24h recordings. **(A1)** Image from the infrared top-view camera utilized for motion tracking. **(A2 & A3)** Representative traces of WT^+/+^ and P120^+/-^ activity over 24h within individual recording chambers. **(B)** Local regressions for distance traveled over 24h by all WT^+/+^, P60^+/-^, and P120^+/-^ mice. **(C)** P120^+/-^ showed significantly higher activity than WT^+/+^ and P60^+/-^ during the exploration and nesting phase (WT^+/+^ vs. P120^+/-^: 1-Way ANOVA; F_2,20_=20.18 P<0.001 p<0.001, P120^+/-^ vs. P60^+/-^: p<0.001). (p<0.05 *, p<0.01 **; p<0.001 ***; post-hoc Bonferroni).

### Activity-dependent theta modulation is impaired during wake in P120^+/-^

Field potential recordings reveal cross–frequency coupling between theta and gamma oscillations^36, 37^. Specifically, gamma oscillations are nested within theta oscillations^38, 39^. In humans, theta-gamma coupling has shown high predictability for correct memory retrieval^40–42^. Mouse studies utilizing optogenetics have demonstrated causal evidence for the importance of theta-gamma coupling in cognitive function^43^. In WT^+/+^ mice, theta and gamma spectral power demonstrated an inverse relationship during wake and sleep states (Supplementary Fig. 5). During wake, the theta-gamma ratio (TGR) was low in WT^+/+^ mice, while during sleep the TGR was high (Supplementary Fig. 5)^44, 45^. This relationship between theta and gamma power was lost in P120^+/-^ as the TGR did not change according to wake or sleep state (Supplementary Fig. 5C - D2). PMP rescued *Syngap1^+/-^* TGR modulation to WT^+/+^ levels.

To determine differences in theta and gamma power during transitions from stationary-wake to active-wake, ten-minute epochs of transitions from stationary- to active-wake (five minutes each) were analyzed (Fig. 6A). During stationary-wake, high theta and low gamma power were apparent in WT^+/+^ mice (Fig. 6A & B). However, during active-wake, WT^+/+^ mice transitioned to high gamma power and low theta power. This state-dependent (stationary- vs. active-wake) transition was not apparent in P60^+/-^ or P120^+/-^ mice (Fig. 6B & C). Specifically, theta power remained consistently high through transitions from stationary- and active-wake in P120^+/-^ mice. Gamma power was not significantly affected during these specific wake-state transitions in P60^+/-^ or P120^+/-^ mice. PMP did not significantly modulate theta during transitions from stationary-wake to active-wake (Fig. 6C).

**Fig. 6.**
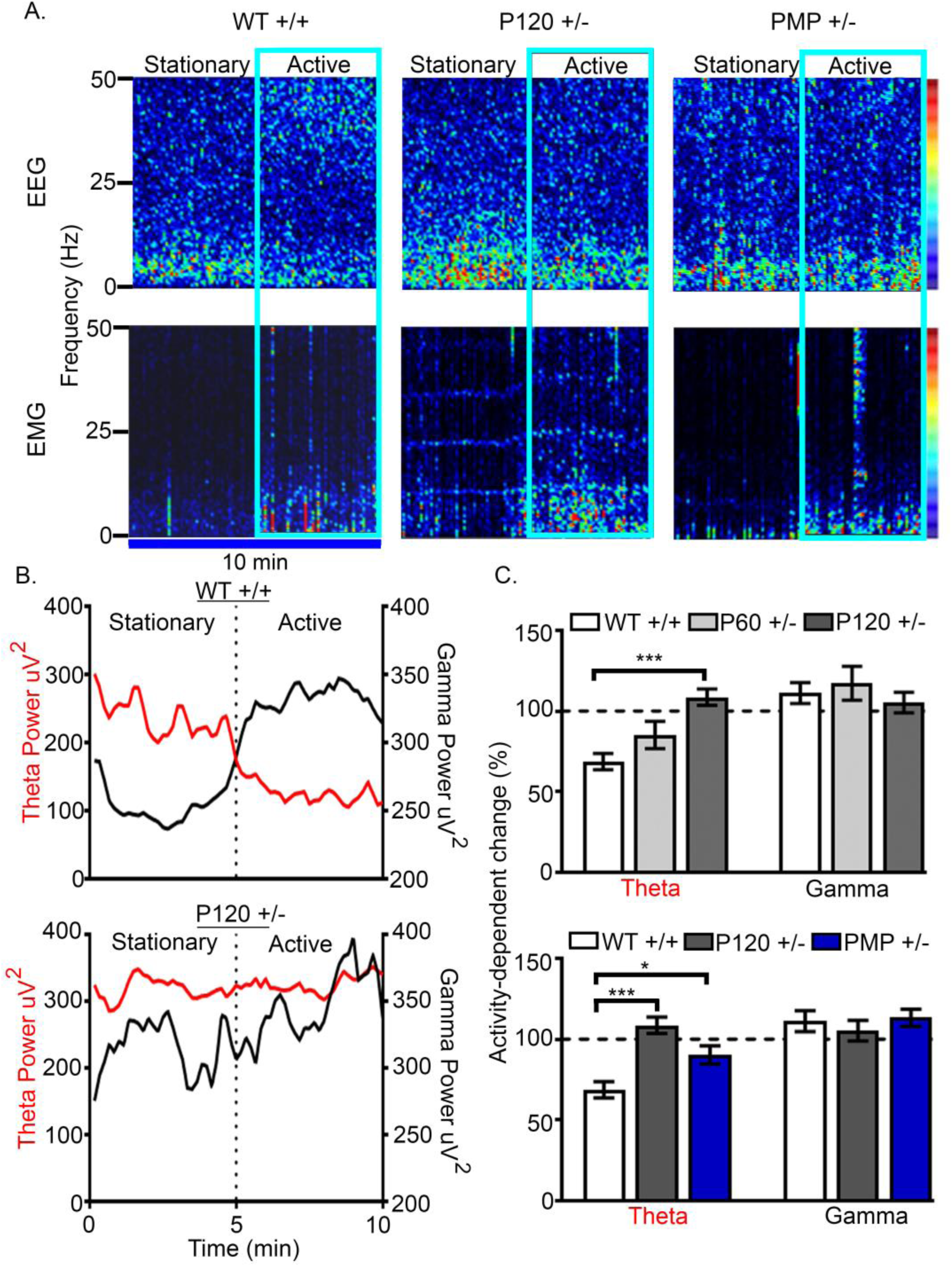
Progressive impairment of theta power during transitions from inactive to active wake. **(A)** EEG and EMG spectral power heat maps of activity-dependent transitions from stationary to active wake in WT mice. P120 ^+/-^ mice failed to demonstrate an increase in gamma with concurrent decrease in theta. **(B)** Representative traces of theta-gamma modulation in WT and P120 ^+/-^ mice during stationary to active transitions. **(C)** The lack of theta modulation during transitions progressively worsened (WT^+/+^ theta vs. P120^+/-^ theta % change: 1-way ANOVA, F_2,19_=12.123, P<0.0001, p<0.0001) PMP administration (2mg/kg; I.P.) at 10am and 6pm improved theta modulation but not to WT levels (WT^+/+^ theta vs. PMP^+/-^ theta % change: 1-way ANOVA, F_2,19_=12.123, P<0.0001, p=0.017). (p<0.05 *, p<0.01 **; p<0.001 ***; post-hoc Bonferroni).

### Loss of behavioral-state homeostasis of cortical gamma

Gamma (35-50Hz) is low during sleep and high during wake^46–48^. High cognitive load during wakefulness^46–48^ is associated with increased gamma power. Accordingly, in WT^+/+^ mice gamma power was high during wake and low during NREM sleep (Fig. 7A). P120^+/-^ mice did not demonstrate behavioral-state dependent gamma power modulation during Wake-NREM transitions, as gamma power remained high during wake and NREM (Fig. 7B). PMP rescued behavioral-state dependent gamma power modulation. PMP-treated *Syngap1^+/-^* mice had high gamma during wake and low gamma during NREM similar to WT^+/+^ mice (Fig. 7C). To analyze differences in Wake-NREM transitions over 24h, slopes for gamma power were calculated for every Wake-NREM and NREM-Wake transition (see Methods). In WT^+/+^ mice, the slope of gamma power during Wake-NREM transitions was negative as gamma power decreased during sleep compared to the previous wake. During NREM-Wake transitions, the slope of gamma power was positive in WT^+/+^ mice as gamma power increased from NREM to wake (Fig. 7D1). The negative slopes of Wake-NREM transitions and the positive slopes of NREM-Wake transitions were significantly different in WT^+/+^ mice (Fig. 7D1). P120^+/-^ mice failed to show significant differences between wake and sleep transitions as gamma power remained high during NREM (Fig. 7D2). PMP rescued the lack of significant differences between the slopes of gamma power during Wake-NREM and NREM-Wake transitions in *Syngap1^+/-^* mice (Fig. 7D3). The average gamma slopes for Wake-NREM and NREM-Wake transitions were calculated to compare groups (Supplemental Fig. 6). Mean gamma slopes for NREM-Wake and Wake-NREM in WT^+/+^ mice were significantly different. In P120^+/-^ mice gamma slopes between NREM-Wake and Wake-NREM were not significantly different. PMP rescued the lack of significant differences between NREM-Wake and Wake-NREM in P120^+/-^ mice (Supplemental Fig. 6).

**Fig. 7.**
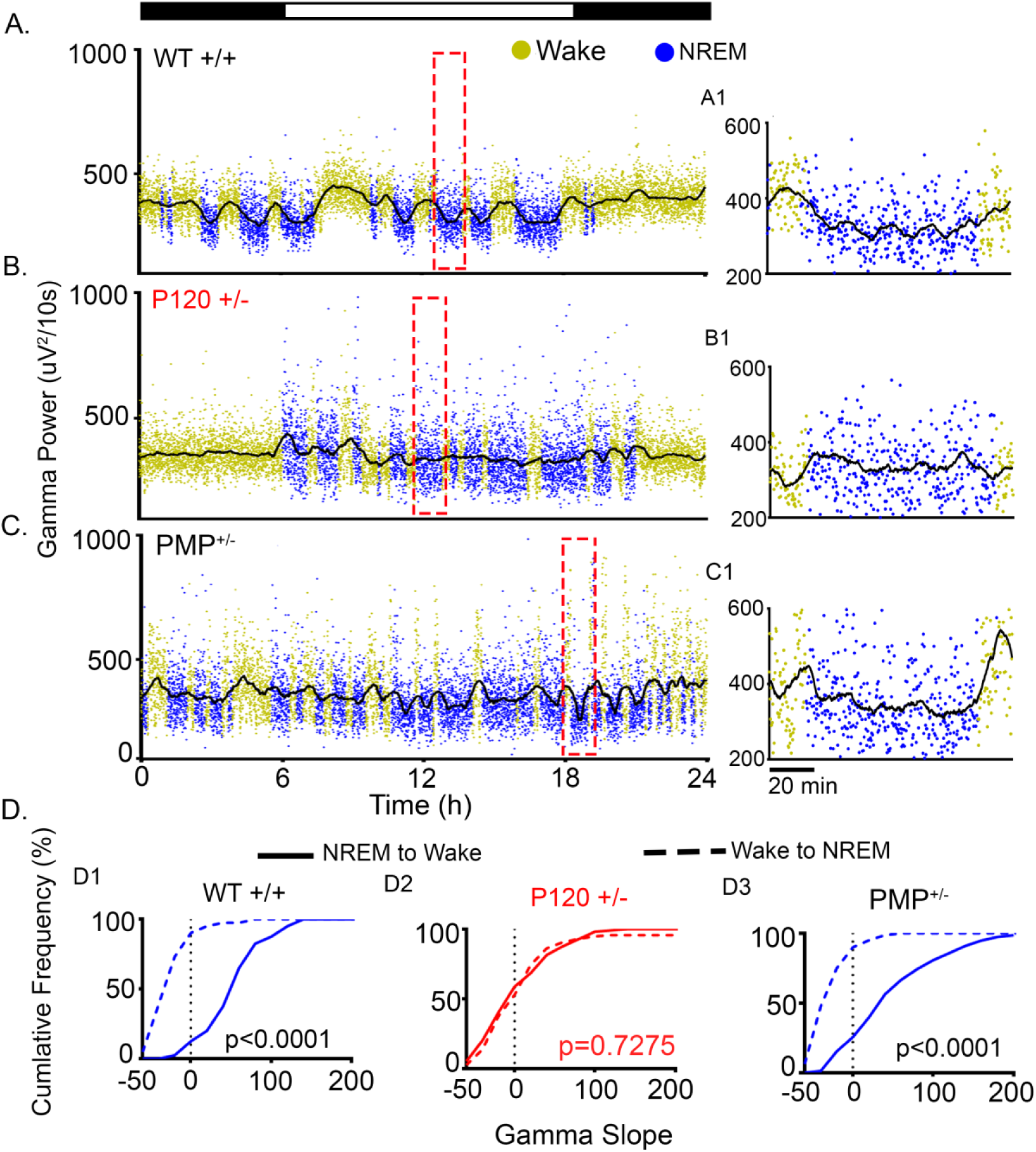
Gamma during transitions between wake and NREM Behavioral state modulation of EEG gamma spectral power. **(A)** 24h WT^+/+^ gamma traces have high gamma during wake and low gamma during NREM. **(A1)** Expanded time scales (red boxes in A-C indicate location of expanded timescales in A1-C1) show a gradual fall of gamma from wake to NREM and gradual rise of gamma from NREM to wake. **(B)** Gamma power in the P120^+/-^ failed to transition to NREM levels, expanded time scale **(B1)** demonstrates the failure of gamma attenuation during a NREM cycle. **(C)** In PMP-administered P120^+/-^ (PMP^+/-^), behavioral-state transitions in gamma power were significantly rescued **(C1)**. **(D)** Cumulative frequency graphs of positive NREM to Wake slopes and negative wake to NREM slopes in WT^+/+^. **(D1)** These slopes were significantly different (*t-test*, t_275_=10.59, p<0.001) between the two independent transition states. **(D2)** No behavioral-state dependent modulation of gamma power occurred in P120^+/-^ (*t-test*, t_327_=0.3488, p=0.7275). **(D3)** PMP administration restored gamma power behavioral-state dependent modulation (*t-test*, t_156_= 9.854, p<0.0001). (p<0.05 *, p<0.01 **; p<0.001 ***).

Frequency analysis of gamma power (35-50Hz) identified differences between NREM and wake in WT^+/+^ mice (Supplemental Fig. 8). Gamma power from 35-45Hz were significantly higher during wake compared to NREM in WT^+/+^ mice. In contrast, in P120^+/-^ mice, wake and NREM gamma power from 35-45Hz was not significantly different (Supplemental Fig. 8). Therefore, *Syngap1^+/-^* mice demonstrated high gamma power during sleep and a loss of state-dependent gamma modulation. Antagonizing AMPARs with PMP rescued state-dependent gamma modulation.

During NREM the majority of brain activity is synchronous slow wave activity with low gamma power^46–48^. In contrast, gamma power increases during the asynchronous activity of REM^46–48^. In WT^+/+^ mice, NREM-REM transitions underwent an increase in gamma, decrease in theta, and decrease in delta power (Fig. 8A & B). P60^+/-^ mice had a significantly greater decrease in delta from NREM to REM compared to WT^+/+^ mice. However, the significant decrease in delta was not apparent in P120^+/-^ mice. In contrast, a significantly lower rate of percent change in gamma power during NREM-REM transitions was apparent in P120^+/-^ mice. Gamma power remained high during both NREM and REM (Fig. 8A & B) in P120^+/-^ mice. The impairment of gamma power modulation from NREM to REM worsened with age, and PMP rescued the modulation of gamma power (Fig. 8C).

**Fig. 8.**
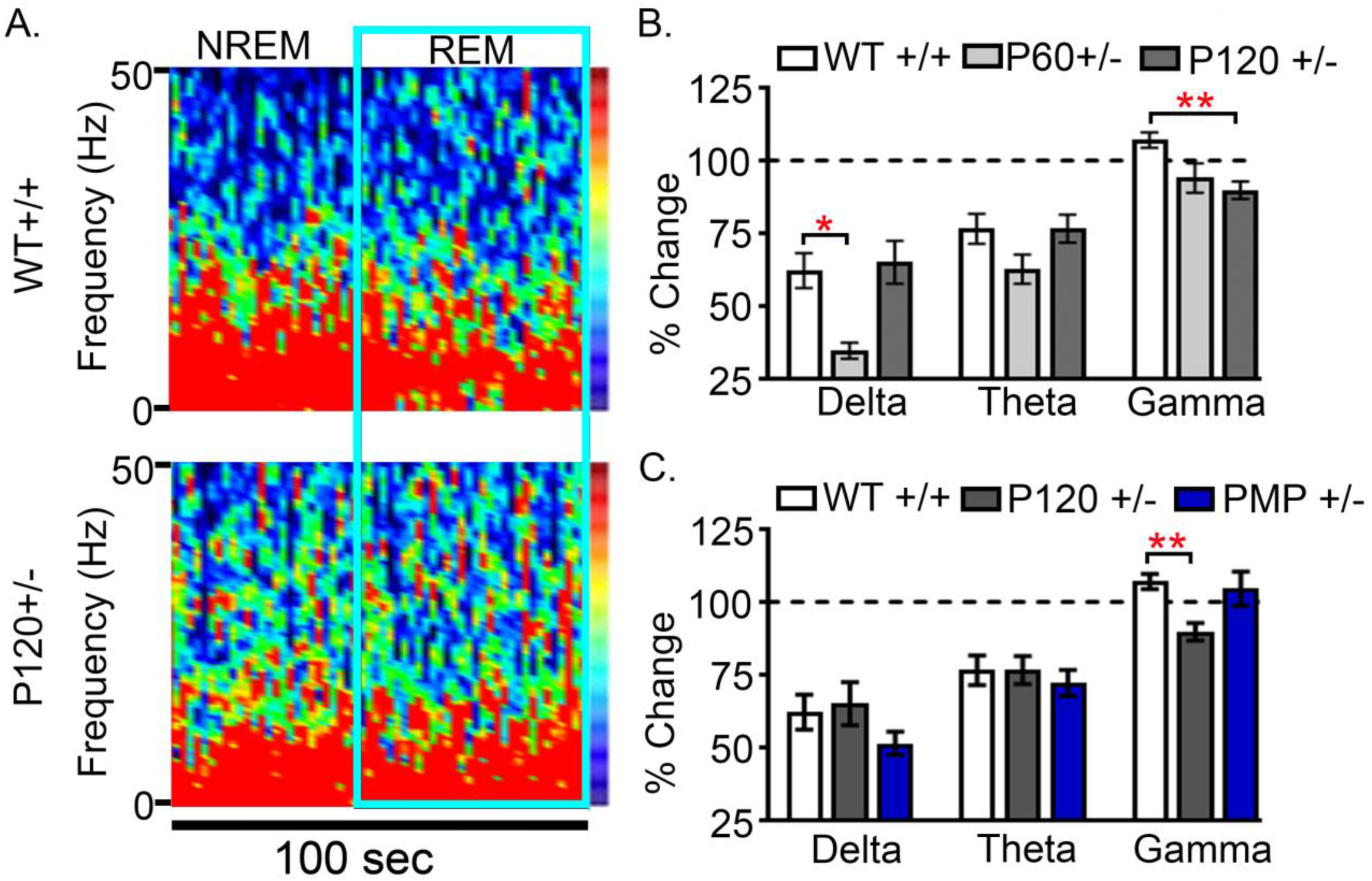
PMP rescued gamma modulation during transitions from NREM to REM. **(A)** Spectral power heat maps demonstrated an increase in gamma power, decrease in theta, and decrease in delta power during NREM to REM transitions in WT ^+/+^mice. **(B)** A decrease in gamma power during NREM to REM transitions was also demonstrated in P120^+/-^ mice. However, delta power during NREM to REM transitions was significantly attenuated in P60^+/-^ mice (WT^+/+^ Delta vs. P60^+/-^ Delta % change: 1-way ANOVA, F_2,97_=2.066, P=0.005, p=0.018) but was absent in P120^+/-^ mice (WT^+/+^ Delta vs. P120^+/-^ Delta % change 1-way ANOVA, F_2,97_=2.066, P=0.005, p=1.00). **(C)** NREM to REM gamma modulation was significantly attenuated at P120^+/-^ (WT^+/+^ Gamma vs. P120^+/-^ Gamma % change 1-way ANOVA, F_2,97_=6.821, P=0.002, p=0.001) but was rescued by PMP administration. (p<0.05 *and p<0.01 **; post-hoc Bonferroni).

### Deficits in PV+IN innervation in prefrontal cortex

The majority of perisomatic inhibition is mediated by parvalbumin-positive interneurons (PV+INs)^46^, whichhave a significant role in cortical circuits and in the modulation of cortical gamma oscillations^49^. The novel finding of consistently high gamma power in *Syngap1^+/-^* mice directed the investigation of their density distribution in the cerebral cortex, hippocampus, and prefrontal cortices. Immunohistochemical (IHC) probes quantifying the PV+IN distributions did not show significant differences in overall densities in the cortex and hippocampus. Layer-specific analyses detected significantly higher numbers of PV+INs in the stratum radiatum (SR) and stratum oriens (SO) regions of hippocampalCA3 and CA1, respectively (*t-test*, t_14_=-2.479, p=0.027, and t_14_=-2.751, p=0.016 respectively). Quantification of overall counts of PV+ puncta identified significant deficits in the dorsal pallidum (DP) cortex in *Syngap1^+/-^* mice (Fig. 9A & B) but not in the prelimbic (PrL) cortex (data not shown). Additionally, a significant decrease in perisomatic PV+ puncta was identified in the DP cortex of *Syngap1^+/-^* mice (Fig. 9B). No significant differences in mean GluA2 fluorescence intensity on PV+INs or non-PV+ neurons were detected in the DP cortex (Fig. 9C1 & 9C2). These findings indicate that deficits in PV+ innervation were region-specific in *Syngap1^+/-^* mice even within the prefrontal cortex.

**Fig. 9.**
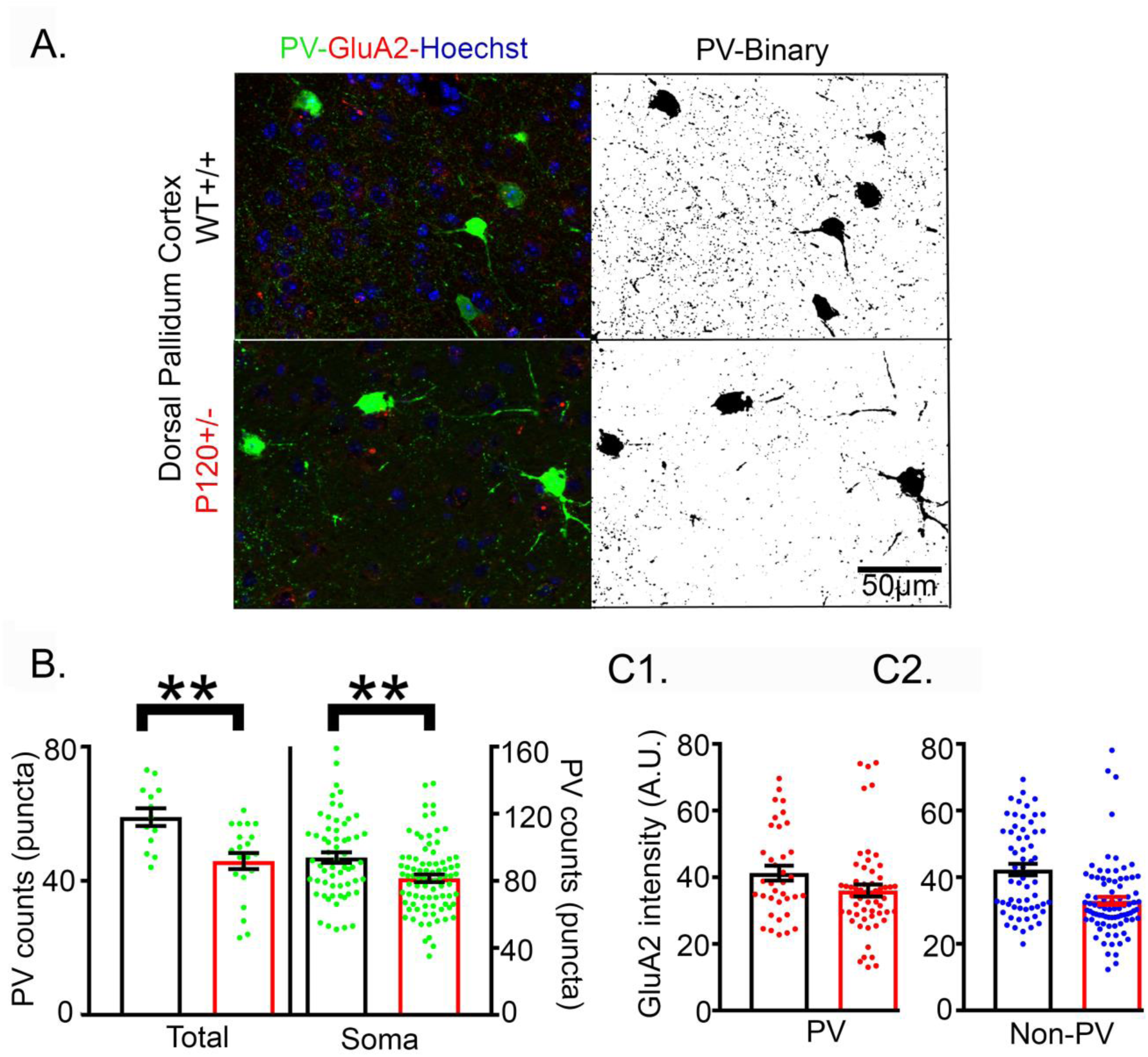
PV+IN deficits in pre-frontal cortex. **(A)** Representative 40X Z-Stack image from the dorsal pallidum (DP) cortex in WT^+/+^ and *Syngap1^+/-^* mice (Het^+/-^) mice. **(B)** PV immunofluorescence represented by PV+ puncta showed a significant reduction in PV innervation in the DP cortex. In Het*^+/-^* mice, total PV puncta counts were significantly lower (*t-test*, t_33_=2.99, p=0.0051). **(C1 & C2)** PV puncta counts onto non-PV somas were also significantly lower in the DP (*t-test*, t_142_=3.25, p=0.0014). There were no significant differences in GluA2 expression on PV+IN or non-PV neurons in the DP cortex. Similar analyses done in the PrL found no significant differences between genotypes. (p<0.05 *, p<0.01 **; p<0.001 ***).

The significant difference between counts of PV+ puncta in the DP cortex (Fig. 9) warranted the analysis of PV+ puncta counts in other regions of interest. Unlike the prefrontal cortex, no significant differences in PV+ puncta count were noted in either the sensory or motor cortices (Fig. 10A2 & B2). A layer-specific investigation of the barrel cortex (BCX) and motor cortex (MCX) demonstrated that PV+ puncta counts were not significantly different between layers 2-3 and 5-6 in the BCX and MCX in both the total counts within the whole acquired images and the somas of the sampled non-PV+ cells (Fig. 10A1, A2 & Fig. 10B1, B2). In the hippocampus, PV+ puncta counts were not significantly different in CA1 or CA3 (Fig. 10C1). Normalized PV+ puncta counts were not significantly different between the SP of WT^+/+^ and P120^+/-^ mice in CA1 and CA3 (data not shown, *t-test*, t_22_=0.2861, p=0.7775, and t_31_=0.8392, p=0.4078 respectively). SO and SR were also examined, and PV+ puncta counts were not significantly different in CA1 and CA3 in the SO (data not shown, *t-test*, t_22_=1.125, p=0.2725, and t_31_=0.5407, p=0.5926 respectively) and the SR (data not shown, *t-test*, t_22_=0.8627, p=0.3976, and t_30_=0.3402, p=0.7361 respectively) between WT^+/+^ and P120^+/-^ mice. Since all *Syngap1^+/-^* mice showed epileptiform activity, the hippocampi were investigated for mossy fiber sprouting. No evidence for mossy fiber sprouting was detected in this model (see Supplemental Fig. 9). In summary, PV+IN deficits in innervation were only observed in the DP cortex.

**Fig. 10.**
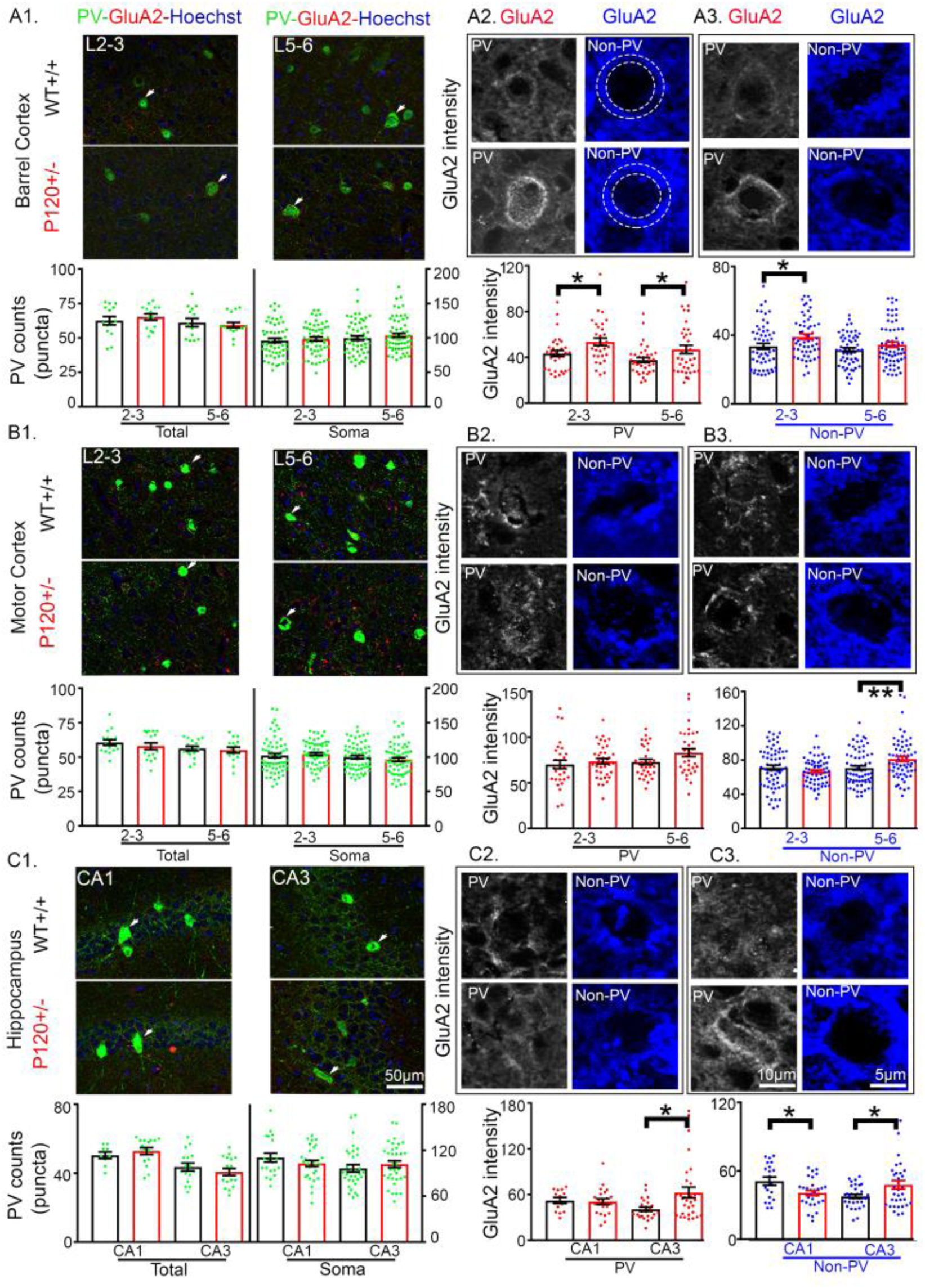
Increased GluA2 expression on PV+ soma were region specific. **(A1)** Representative 40X Z-stack images of WT^+/+^ and *Syngap1^+/-^* mice (Het^+/-^) from barrel cortex (BCX) layers 2-3 and 5-6. White arrow heads in A1 indicate PV+IN shown in A2 and A3 for BCX layers 2-3 and 5-6, respectively. Total PV puncta counts and somatic PV puncta counts were not significantly different in the BCX. Representative 40X Z-stack images of GluA2 immunofluorescence on PV+IN and non-PV neurons for layers 2-3 and 5-6 **(A2 & A3)**. Het^+/-^ GluA2 expression was significantly higher on PV+IN in both BCX layers 2-3 (*t-test*, t_63_=2.437 p=0.0176) and 5-6 (*t-test*, t_70_=2.219, p=0.0297). Het^+/-^ non-PV neurons in BCX layer 2-3 also had a significant increase in GluA2 expression (*t-test*, t_110_=2.405, p=0.0178). **(B1)** Representative 40X Z-stack images from motor cortex (MCX) layers 2-3 and 5-6, total PV puncta counts and somatic PV puncta counts were not significantly different in the MCX. White arrow heads in B1 indicate PV+IN shown in B2 and B3 for MCX layers 2-3 and 5-6, respectively. **(B2 & B3)** GluA2 expression was not significantly different for PV+IN in the MCX layers 2/3 and 5/6. Het^+/-^ GluA2 expression was significantly increased in PV+IN of MCX layers 5/6 (*t-test*, t_126_=2.707, p=0.0077). **(C1)** Representative 40X Z-stack images from hippocampal CA1 and CA3. White arrow heads in C1 indicate PV+IN show in C2 and C3 for CA1 and CA3, respectively. Total PV puncta counts and somatic PV puncta counts were not significantly different in neither CA1 nor CA3. Representative 40X Z-stack images of GluA2 immunofluorescence on PV+IN **(C2)** and non-PV neurons **(C3)**. CA3 Het^+/-^ PV+IN had an increase in GluA2 expression (*t-test*, t_52_=2.622, p=0.0114; C2). In non-PV neurons, CA3 GluA2 expression was also increased in Het^+/-^ mice (*t-test*, t_64_=2.428, p=0.018; C3). In contrast, CA1 Het^+/-^ non-PV neurons expressed significantly less GluA2 (*t-test*, t_48_=2.582, p=0.0129; C3). (p<0.05 *, p<0.01 **; p<0.001 ***).

### Increased GluA2 expression on PV+ soma in somatosensory but not motor cortex

A rich diversity of GABAergic neurons shape the spatiotemporal dynamics of cortical circuit outputs by elegant inhibitory control mechanisms^50, 51^. Identification of the dysregulated cortical gamma and its acute rescue by PMP directed the quantification of AMPA-GluA2 on PV+INs and non-PV+ neurons in these regions. Compared to neurons, PV+INs are known to have a lower expression profile for the GluA2 subunit^52^. The findings (Fig. 10) highlighted a region-specific alteration for the significant upregulation of GluA2 in both PV+INs and non-PV+ neurons (for GluA2 antibody verification see Supplemental Fig. 9). The sensory and motor cortices in the *Syngap1^+/-^* mice were distinctly different. PV+INs in the both 2-3 and 5-6 cortical layers of the BCX showed significant upregulation, (Fig. 10A2) which was absent in the MCX (Fig. 10B2). Significant GluA2 upregulation in non-PV+ sampled cells in both ROIs was evident in layer 2-3 of the BCX and layer 5-6 of the MCX (Fig. 10B2). The mean GluA2 fluorescence intensity on PV+INs was also significantly higher in the CA3 but not in the CA1 (Fig. 10C2). For non-PV+ somas, the mean GluA2 fluorescence intensity was significantly lower in the CA1 but significantly higher in the CA3 neurons (Fig. 10C2). Therefore, the results identified a significant dysregulation of GluA2 expression in PV+INs, a population of INs known to contribute significantly in cortical gamma oscillations^49^. The regional differences highlight the complexity of the outcomes *Syngap1^+/-^* haploinsufficiency can exert in different circuits.

Additionally, a higher mean GluA2 fluorescence intensity was observed in the MCX compared to the BCX. In WT^+/+^ mice, the mean GluA2 fluorescence intensity in MCX neurons was significantly higher than BCX neurons both for layers 2/3 and 5/6 (Fig. 10A2 vs. 10B2, *t-test*, t_117_=10.6, p<0.0001, and t_118_=12.94, p<0.0001 respectively). This relationship was preserved in P120^+/-^ mice, as the mean GluA2 fluorescence intensity in MCX neurons was also significantly higher than in BCX neurons in both layers 2/3 and 5/6 (Fig 10A2 vs. 10B2, *t-test*, t_119_=11.31, p<0.0001, and t_126_=14.18, p<0.0001 respectively).

## Discussion

In this study, we used long-term EEG monitoring and qEEG protocols developed specifically to identify biomarkers underlying epileptogenesis in mouse models of acquired and genetic NDDs^17, 18, 53^. The results showed a progressive worsening in epileptogenesis associated with significant alterations in macro- and micro-sleep architecture, IIS frequency, and impaired behavioral-state dependent cortical activity during wake- and sleep-state transitions. The IISs and seizures in the *Syngap1^+/-^* mice predominantly arose during NREM sleep. In the human overnight EEG analysis, identification of ictal clusters during NREM were a novel finding that support the translational validity of this *Syngap1^+/-^* mouse model. These results highlight the similarities in seizure phenotypes in this mouse model to phenotypes described in the clinic. Seizures are highly prevalent in patients with MRD5, therefore these findings provide a valuable contribution to help further understand the significance of *Syngap1* in epileptogenesis. The intellectual disability and behavioral abnormalities i in *Syngap1* heterozygous mice have been well described^34, 54^. However, the seizures in *Syngap1* heterozygous mice have only been identified and not thoroughly characterized until now. Here we report the first thorough analysis of the epilepsy in a model of *Syngap1* haploinsufficiency. Currently, it is unknown if other models of *Syngap1* haploinsufficiency that differ by mutation site or mutation strategy (i.e. germ-line vs. Cre-recombinase) have recurrent spontaneous seizures.

Spectral power analysis of the mouse EEGs quantified the cortical gamma homeostasis during behavioral-state transitions and identified a significant disruption in *Syngap1^+/-^* mice. The rescue of this biomarker with acute dosing by PMP identified a critical role of AMPARs in gamma homeostasis during such transitions. Given the novel finding of acute rescue of gamma homeostasis by an AMPAR antagonist PMP and previous work implicating severe disruption of gamma oscillations associated with induced increase of GluA2 expression in INs^55^; the Ca^2+^ impermeable AMPA subunit became a primary focus. This approach identified excessive GluA2-AMPAR expression on the somas of PV+INs that was location- and circuit-specific indicating a significant role of PV+INs in MRD5 pathology. This study also found abnormal cortical activity in theta frequencies during transitions from stationary-wake to active-wake. As theta-gamma coupling is essential for memory retrieval, abnormal theta and gamma homeostasis pose a potential relevant biomarker for memory impairment that has been previously verified in human subjects and optogenetic rodent studies^40–43^.

To date, there is no clear correlation between *SYNGAP1* mutation location and phenotypic severity. However, mutations in exons 8-15 seem to be associated with seizures more pharmacoresistant than with mutations in exons 4-5^11^. In this preclinical model, deletions in *Syngap1* exons 8 and 9 resulted in half of the *Syngap1^+/-^* mice presenting with a seizure during the 24 recording period indicating a high seizure frequency. However, 100% of the *Syngap1^+/-^* mice presented progressively increasing IIS frequency and sleep dysfunction. These findings provide a potential mechanism for the reported decline of cognitive ability during the developmental trajectory of MRD5 into adulthood^21^.

Wakefulness is associated with net-synaptic potentiation, whereas sleep favors global synaptic depression thereby preserving an overall balance of synaptic strength^56^. AMPAR levels are high during wakefulness and low during sleep. Therefore, consistently high AMPAR levels in PV+INs would perturb this balance toward impaired signal processing and learning disability. qEEG analyses revealed significant loss in cortical gamma homeostasis associated with behavioral-state transitions both in wake and sleep that could impede cortical information processing. Gamma oscillations^57, 58^ are known to heavily depend upon AMPAR kinetics in interneurons, especially for long-range synchrony. We showed that acute treatment with PMP, an AMPAR antagonist, significantly rescued many of the identified EEG biomarkers along with the disrupted cortical gamma oscillations. IHC analyses observed and quantified a significant increase in GluA2-AMPAR expression onto the somas of cortical and hippocampal PV+INs in a region-specific manner. The significant rescue of the cortical gamma homeostasis by acute dosing of low-dose PMP when compared to the weaker effect on IIS suppression may indicate a novel role for PMP unrelated to its FDA approved anti-seizure role. New trials evaluating PMP efficacy in infants^23^ are underway. As an anti-seizure agent for pediatric epilepsies, PMP dosing is gradually ramped-up starting with the low-dose of 2mg/kg with weekly increments reaching maintenance doses of 8-12 mg/day based on tolerability. That is when its anti-seizure efficacy is evaluated. This study showed that low-dose PMP has a novel and acute role in rescue of gamma homeostasis in *Syngap1^+/-^* mice likely unrelated to the maintenance dosing requirement for its anti-seizure efficacy. PMP anti-seizure efficacy for SYNGAP1 are currently not known and a recent cohort study with 57 patients reported only one patient having received PMP treatment. The findings reported here indicate the necessity to develop GluA2-AMPAR selective antagonists, which currently do not exist. These analyses also identified poor PV+ interneuron innervation in the prefrontal cortex that was not detected in the other ROIs evaluated both in the cortex and hippocampus. This finding in the prefrontal cortex is supported by previously published reports^59^.

Increasingly, brain oscillations are being used to understand complex neuropsychiatric disorders. Gamma (35–50Hz) oscillations have warranted special attention due to their association with higher order cognitive processes including sensory processing, attention, working memory, and executive functioning. Activation of GABAergic interneuron networks has been shown to produce gamma oscillations (∼40Hz) in both the hippocampal and neocortical networks. Given the critical role of PV+INs in cortical gamma oscillations^49, 60^ and the role of AMPARs in homeostatic synaptic plasticity^61^, the loss of gamma homeostasis identified for *Syngap1* haploinsufficiency during periods commonly associated with intense synaptic plasticity^62^ provide novel insights into the associated intellectual disabilities.

GluA2 subunits confer Ca^2+^ impermeability to the AMPAR, a slow EPSP in excitatory neurons, and are expressed at low levels in GABAergic interneurons^57^. AMPA-mediated currents rise and decay faster in interneurons partly because of these differences in subunit profiles^63, 64^. A genetically altered GAD-GluR-B (GluA2) mouse expressing high levels of the GluA2 subunit in GAD+ cells exhibited a severe disruption in gamma oscillations between spatially separated sites and significant changes in interneurons firing patterns^57^. PV+INs receive convergent excitatory input from principal neurons, and inhibitory input primarily from other PV+INs. Experiments that have causally tested stimulation of PV+ cells *in vivo* in the BCX by ontogenetic manipulation selectively amplified gamma oscillations^65^. Importantly, this activation suppressed the sensory processing in nearby excitatory neurons within the BCX. Findings of increased GluA2 AMPARs on PV+ soma both in layer 2/3 and 5/6 of the BCX would predict similar suppression of sensory processing in the BCX. Findings supporting this hypothesis have recently been documented in the BCX of a mouse model of *Syngap1* haploinsufficiency^66^. GluA2 data for PV+INs in the BCX but not in the MCX, highlight the divergent circuit- and location-specific effects of *Syngap1* haploinsufficiency. Similarly significantly higher mean GluA2 fluorescence intensity in layer 5-6 motor output neurons in the MCX highlight location-specific effects for non-PV+ neurons. These data also indicate that PV+IN hyperexcitability or lack thereof may be one of the underlying factors that determine circuit outcomes to motor vs. sensory inputs. An increase of GluA2 expression in PV+INs in the BCX could implicate recruitment of excessive PV+IN mediated inhibition resulting in cortical hyperexcitability via rebound excitation. This hypothesis may underlie the clinical reports of reflex seizures in MRD5 emerging during activities associated with increased cortical gamma synchronicity^12, 28^.

Morphology of PV+INs has been identified as unipolar vs. multipolar^67^ and little is known about the differences in AMPA-mediated currents in these subtypes. GluA2-lacking AMPARs promote anti-Hebbian long-term plasticity^68^ which is critical for projection interneurons that functionally connect spatially distant circuits during development. Increase of GluA2-AMPARs in PV+IN soma could be one novel mechanism underlying autistic-like behaviors, ID, and seizures in *SYNGAP1* haploinsufficiency. Little is known of the developmental and regional expression profiles of SYNGAP1 isoforms, however it is known is that they can exert opposing effects on synaptic strength^69^. Therefore, region-specific and isoform-specific haploinsufficiency could confer vastly disparate and sometimes opposing circuit-specific outcomes within the same brain. The findings of this study and a previous report^70^ support this conclusion.

Synaptic strength is an important factor in learning and memory that is regulated by complex biochemical machinery at the PSD. There are currently no evidence-based clinical interventions that can directly modulate SYNGAP1 function. One of the prominent roles of SYNGAP1 is to regulate synaptic plasticity, and this process is heavily involved in both epileptogenesis and sleep homeostasis. Therefore, using validated qEEG biomarkers for both epileptogenesis and circuit dysfunction in preclinical models could be a sound translational approach. Investigation of circuit function in intact brains as they transition from active exploratory, inactive-wake, and REM/NREM sleep states is an exciting frontier. These are behavioral-state transitions that are most reliant upon synaptic plasticity.

## Methods

All animal care and procedures were carried out in accordance with the recommendations in the Guide for Care and Use of Laboratory Animals of the National Institutes of Health. All protocols used in this study were approved by the Committee on the Ethics of Animals Experiments of the Johns Hopkins University. In *Syngap1*^+/-^ mice, exons 7 and 8 (C2 domain) were replaced with a neomycin resistance gene in the opposite direction to eliminate all possible slice variants^26^. All mice were single-housed at P46 (1 week before electrode implantation surgeries) in polycarbonate cages with food and water provided *ad libitum*, on a 12h light-dark cycle. All efforts were made to minimize animal suffering and the total number of animals used.

### EEG surgery and 24h video-EEG/EMG recording

Surgical procedures implemented in this study are previously described^71^. All surgical procedures were performed under isoflurane anesthesia (4%-2.5%). 7 days (±1 day) before recording, mice were surgically implanted with subdural EEG electrodes and suprascapular EMG electrodes (Fig. 1A & C). EEG electrodes utilized coordinates from bregma for consistent placement^18^.

After recovering from electrode implantation surgery (7±1 day) mice were placed in a recording chamber with food and water provided *ad libitum*. WT^+/+^ and *Syngap1*^+/-^ mice underwent 24h vEEG/EMG at younger (P60) and older (P90-120) ages (see Supplementary Fig. 1 for sample size). All 24h vEEG/EMG were recorded using Sirenia Acquisition software (Pinnacle Technology Inc., Lawrence, KS, USA). Each chamber had a top-view infrared camera and a tethered pre-amplifier connected to a commutator. Connecting the commutator to the implanted head cap allowed for unrestricted movement within the recording chamber during recording. EEG recordings were acquired at 400Hz with a 100 gain, band passed from 0.5 to 50Hz^72^. Before each recording, all mice received a new piece of nesting material and a small piece of nesting from their home-cage. Data from all wild type recordings at P60, 90 and 120 were pooled together (see Supplementary Fig. 4) for analysis, P90 *Syngap1*^+/-^ (small sample size; n=2) recordings were pooled with P120^+/-^ mouse recordings. At P121, half of the mice in the study underwent another 24h recording after being administered low-dose 2mg/kg PMP (in (100% EtOH; Aobious) intraperitoneal at 10am and again at 6pm in the 24h recording period.

### Hypnogram and EEG power analysis

EEG data were manually scored for wake, NREM, and REM EEG states in 10s epochs by a scorer blinded to genotype and sex. All EEG artifacts were identified after EEG and video confirmation. Spectral power data for these identified epochs was excluded from further analyses to avoid contamination of data from background EEG. qEEG spectral analysis utilized delta (0.5-4.0Hz), theta (5.5-8.0Hz), alpha (8.0-13.0Hz), beta (13.0-30Hz), and gamma (35-50Hz) frequencies. The data underwent fast Fourier transform (FFT) and were exported for all respective frequencies from an automated analysis module within Sirenia Sleep Pro (Pinnacle Technology Inc., Lawrence, KS, USA).

### Seizure and IIS scoring

Spontaneous seizures were identified by manual review of all 24h EEGs. Identified EEG seizure activity was then further classified by phenotype (review of synchronous video and EMG) and duration. EEG epochs containing seizure activity were also excluded from spectral power analyses of background EEG and spike frequency analysis. IISs were visually analyzed on raw EEG using conservative parameters obtained from obvious pre- and post-ictal spikes within the same dataset. Each subject’s mean EEG amplitude during wake was used to determine the minimum absolute value for IIS identification. 1X of the absolute value was the criterion for the positive inflection, and 2X of the absolute value was the criterion for the negative inflection. If the positive and negative inflection criteria were met, the synchronous video was analyzed to eliminate artifact contamination (e.g. scratching, grooming or drinking water). IISs were manually scored for every 5s EEG epoch to enhance the resolution of IIS frequency detection.

### Gamma AUC and slope

Spectral power data for each mouse was exported for area under the curve (AUC) analysis using trapezoidal summations^53, 71^. FFT generated spectral power data for each mouse underwent AUC analysis in R statistical software with automated code.

To analyze differences in gamma power transitions during changes in behavioral states (i.e.; from sleep to wake and wake to sleep), only behavioral states that lasted ≥6min were used. Two one-minute averages of gamma power AUC were calculated, the first occurring right before the behavioral-state transition and the second occurring 5min after the behavioral-state transition. The slope between these two gamma power averages was then calculated by automated code for each behavioral-state transition in R statistical software^71^.

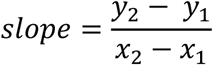

Where y_2_ = gamma power average after transition, y_1_ = gamma power average before transition, x_2_ = 6min, and x_1_ = 1min.

### Theta-gamma ratio and activity dependent theta-gamma modulation

Theta-Gamma Ratio (TGR) was calculated for every 10s epoch for the 24h recording period. To analyze differences between wake and sleep TGR changes between WT^+/+^ and *Syngap1*^+/-^ mice, the percent change in TGR from wake to sleep was calculated. Wake activity dependent theta-gamma modulation was analyzed by pulling transitions of equal duration (10min total) from stationary-wake to active-wake (5 min each; verified by EMG and video). Respective powers were summed according to their activity state, and then computed for percent change. NREM-REM dependent theta-gamma modulation was analyzed by examining NREM-REM transitions of equal duration (100s total). Respective powers were summed according to their sleep state and percent change was calculated.

### Analysis of human EEG recording

Continuous overnight EEGs are being acquired with parent consent under approved IRB protocol^30^. The raw EEG recording analyzed here was imported into Brainstorm (Brainstorm, CA, USA). The EEG channels F4, C4 and O2 were re-referenced to A1 or A2 and exported in the European Data File (EDF) format, de-identified and assigned an ID number. A bandpass filter of 0-80Hz was used. Automated spectral analysis was conducted (delta 0.5–4Hz, theta 5.5–8.5Hz, alpha 8–13Hz, beta 13–30Hz and gamma 35–80Hz) using the Sirenia Sleep score module and spectral powers were calculated for every 10s epoch of the EEG. Each epoch was manually scored as Wake or Sleep (NREM or REM) using video and delta power during slow-wave sleep in NREM using a previously published protocol^17^. These analyses were performed blind to the time points of the ictal events on the raw EEG. Ictal events were identified on original raw EEG recorded as a 24-channel digital video EEG using an international 10-20 system of electrode placement. Eye monitors and one channel EKG recordings were also available for review. The raster plot for identified ictal events on this 20h long recording was then superimposed over the hypnogram generated for the same EEG recording.

### PV and GluA2 immunofluorescence acquisition and quantification

At P127 (P120+ 7 days) mice were anesthetized with chloral hydrate (300mg/ml; IP) before transcardiac perfusion with ice-cold PBS followed by formalin (10%). Brains were then cryoprotected and stored at −80°C before cryosectioning. All coronal sections were cut at 14um and mounted onto Permafrost™ microscope slides. The primary antibodies used were anti-GluA2 (ThermoFisher Scientific; RRID: AB_2533058; diluted 1:100), anti-parvalbumin (Swant; RRID: AB_2313848; diluted 1:5000), and anti-ZnT3 (Synaptic Systems; RRID: AB_2189664; diluted 1:100). Hoechst 33342 fluorescent dye (ThermoFisher Scientific; Cat. H3570) was used to label cell nuclei. The secondary antibodies used were: donkey anti-rabbit Alexa 488 (ThermoFisher Scientific; RRID: AB_2535792; diluted 1:500), and goat anti-mouse Alexa 594 (ThermoFisher Scientific; RRID: AB_2556549; diluted 1:500). For GluA2 antibody verification, P60 GluA2 KO mice (Richard Huganir, Johns Hopkins University) and age matched littermates were perfused with PBS and 4% PFA then sectioned at 50μm (Supplemental Fig. 9).

Fluorescent images were acquired with Axiovision Software 4.6 and Apotome® (Carl Zeiss, Jena, Germany). Images from all regions of interest (ROIs): hippocampal regions CA1 and CA3, MCX, BCX, and prefrontal cortex regions DP and PrL, were acquired at both 10X and 40X magnifications for all ROIs. The 10X images were acquired as Z-stack mosaics of four 3μm steps, while 40X images were acquired as Z-stack mosaics of four 1μm steps. For 10X and 40X images, two images per animal were analyzed from each ROI. All ZnT3 and GluA2 KO stained hippocampi were imaged as 10X mosaics same as above. All data analyses were performed using the ImageJ software^73^. All data were acquired and analyzed by a scorer blinded to both genotype and sex.

All Z-stack images were fused with equal number of step sizes (four 3μm steps for 10X images and four 1μm steps for 40X images) and split by RGB channel in ImageJ. PV+ Alexa 488 immunofluorescence identified both PV+IN cell bodies and PV+ puncta representing their presynaptic terminals innervating non-PV cells in the same ROIs. PV+ puncta around two randomly selected non-PV+ cells per ROI were analyzed. In this study, non-PV+ cells represented neurons that were selected by the large size of their nuclei with multiple PV+ puncta representing PV innervation. A low-pass filter was used to exclude non-specific labeling of structures larger than 10 pixels (such as blood vessels and RBC), and a high-pass filter was used to exclude structures smaller than 2 pixels (such as nonspecific fluoropore staining on the surface of the brain section). The filtered PV immunofluorescence was then converted to a binary image and PV+ puncta counts were quantified using ImageJ. Total PV+ puncta counts were quantified using ImageJ. For the stratum oriens (SO), stratum pyramidale (SP), and stratum radiatum (SR) layers were delineated and quantified individually for hippocampal images. PV+ puncta were expected to be higher in regions of greater neuron density, such as the SP layer in the CA1 and CA3. Therefore, the PV+ puncta counts were additionally normalized by each sub-ROI area of SP, SO, and SR for every hippocampal image. The somas of non-PV+ cells were circumscribed using an elliptical selection tool to demarcate the neuronal cell body, and puncta immediately around the soma were manually counted. Counts of PV+ puncta on non-PV+ neurons were normalized to their respective cell areas.

To quantify GluA2 expression on PV+INs, mean somatic GluA2 fluorescence (Alexa 594) intensity was quantified for all PV+INs within each ROI. PV+INs were sub-classified by their location in layers 2/3 vs. 5/6 in MCX and BCX images. In hippocampal images, PV+INs were classified by their location in CA1 vs. CA3 and by layer specificity in the SO, SP, or SR. Two randomly selected non-PV+ cells were quantified per ROI in the MCX and BCX. In hippocampal images, two non-PV+ cells were quantified only for the SP. Mean somatic GluA2 fluorescence intensity was quantified as the average pixel intensity (0-255) around each cell body.

### Statistical analyses

Statistical tests were performed using SPSS24 (IBM, Armonk, NY, USA) and Prism 7.0 (GraphPad Software, La Jolla, CA, USA). All 1-way and 2-way ANOVAs were performed with Bonferroni’s post-hoc corrections. Independent *t-*tests were two-way, and all data reported as means were ± 1 S.E.M. Analysis of seizure durations between P60^+/-^ and P120^+/-^ was an independent two-way *t-*test in Prism 7.0. Analyses of macro and micro-sleep architecture were 2-way ANOVAs with Bonferroni post-hoc corrections in Prism 7.0. IIS frequency and motor hyperactivity analyses were 1-way ANOVAs with Bonferroni post-hoc corrections. The running weighted averages of IIS frequency and hyperactivity were plotted using the Lowess method in Prism 7.0.

Theta power during transitions from inactive to active wake were analyzed as 1-way ANOVAs and representative traces of theta-gamma modulation were plotted using the Lowess method in Prism 7.0. Representative TGRs underwent a min-max normalization and were plotted in Prism 7.0. Average % increase in TGR from wake to sleep were analyzed as a 1-way ANOVA with Bonferroni post-hoc correction in Prism 7.0.

Differences within NREM-wake and wake-NREM gamma slopes for each group were analyzed as independent two-way *t-*test in Prism 7.0. To identify differences between NREM-wake and wake-NREM between groups, the average gamma slopes for each transition state were analyzed as a 2-way ANOVA with Bonferroni post-hoc correction in Prism 7.0. Gamma frequency plots of Wake, REM, and NREM were plotted using the Lowess method in Prism 7.0. Wake and NREM spectral frequency were analyzed as two-way independent *t-*tests in Prism 7.0.

Mean PV+IN puncta counts and mean GluA2 florescence intensity from 40X Z-stack images were analyzed in Prism 7.0 as two-way independent *t-*tests.

## Acknowledgements

We thank the families who have consented to participate in the post-hoc analyses of overnight EEGs recorded in their children with identified pathogenic *SYNGAP1* mutations. The authors would like to thank Dr. Jonathan Pevsner for his thoughtful comments and suggestions. Research reported in this publication was supported by the Eunice Kennedy Shriver National Institute of Child Health and Human Development of the National Institutes of Health under Award Number R01HD090884 (SDK). The content is solely the responsibility of the authors and does not necessarily represent the official views of the National Institutes of Health.

## Author Contributions

S.D.K. designed the experiments. S.D.K and R.L.H conceptualized the project. S.D.K. and B.J.S performed experiments. Y.A. managed mouse colonies. B.J.S., S.A., and P.A.K. analyzed data. S.D.K. and B.J.S interpreted the data. S.D.K and B.J.S wrote the manuscript. S.D.K., B.J.S., and P.A.K. edited and reviewed the manuscript.

## Corresponding author

Correspondence to S. D. Kadam.

---

**Supplemental Fig. 1.**
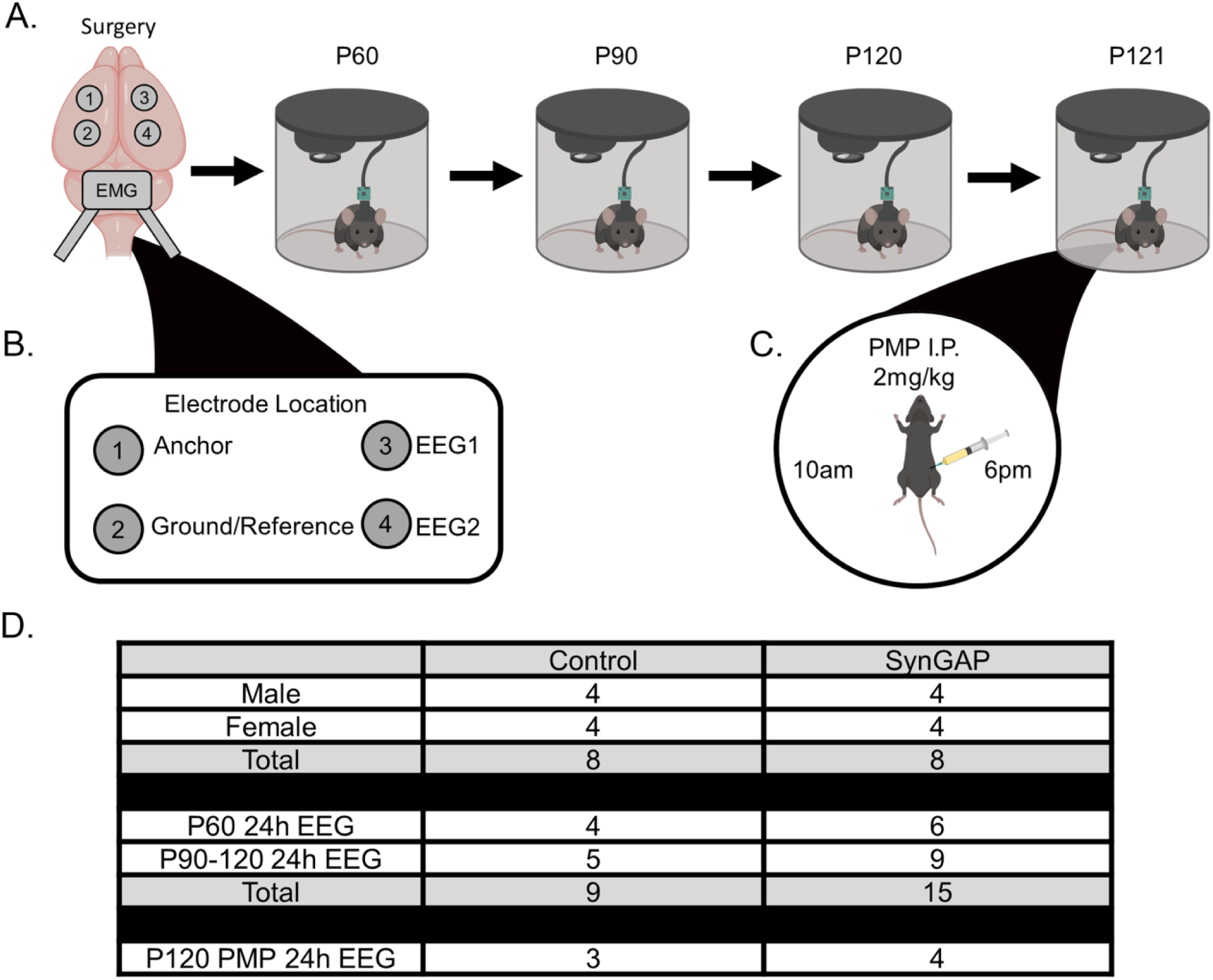
Experimental paradigm to evaluate epilepsy in *Syngap*1^+/-^ mice. **(A)** *In vivo* EEG recordings of mice at P60, P90, P120, and P121. **(B)** At P121 7 out of 16 mice received low-dose Perampanel (2mg/kg I.P.) at 10am and 6pm during the additional 24h recording period. (Schematic generated with help of Biorender) **(C)** Location of electrode placement. **(D)** Table for sample sizes of mice in the study and samples sizes of all the 24h recordings analyzed in this study.

**Supplemental Fig. 2.**
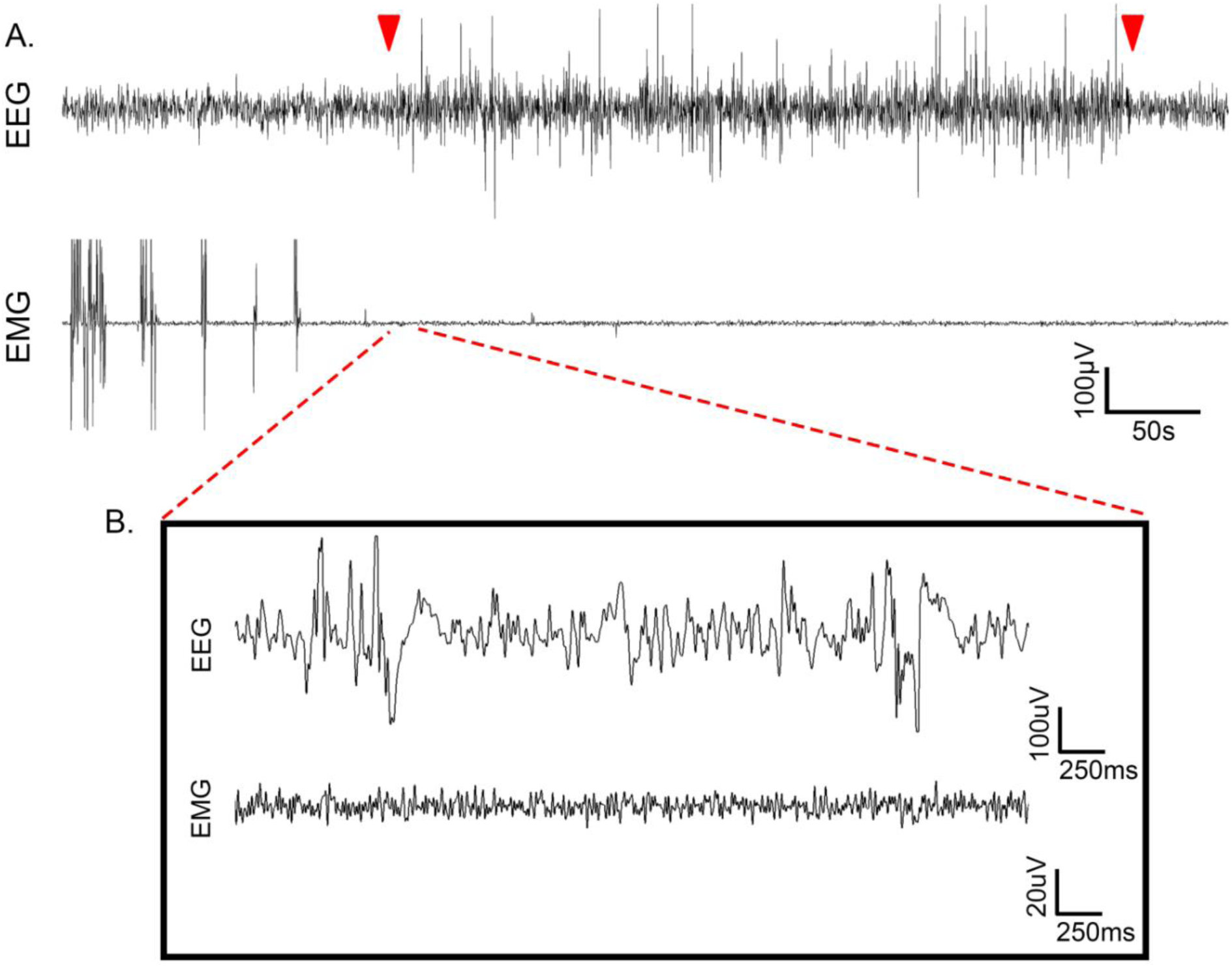
Spontaneous electrographic seizures during wake in *Syngap1*^+/-^ mice. **(A)** A 10min representative EEG trace (0.5-50Hz) of a spontaneous electrographic seizure during wake in a P120^+/-^ mouse (red arrows denote the start and end of the electrographic seizure). Synchronous EMG recording demonstrated attenuation of EMG activity associated with freezing behavior in the mouse noted on video. This phenotype was in contrast to short duration myoclonic seizures (Fig. 1) with ∼3Hz spike-wave discharges on EEG that showed time-locked myoclonic jerks. All electrographic seizures emerged during wake, whereas myoclonic seizures predominantly emerged during NREM. **(B)** Expanded time scale showed ictal activity on EEG recordings with concurrent silent EMG during the freezing behavior.

**Supplemental Fig. 3.**
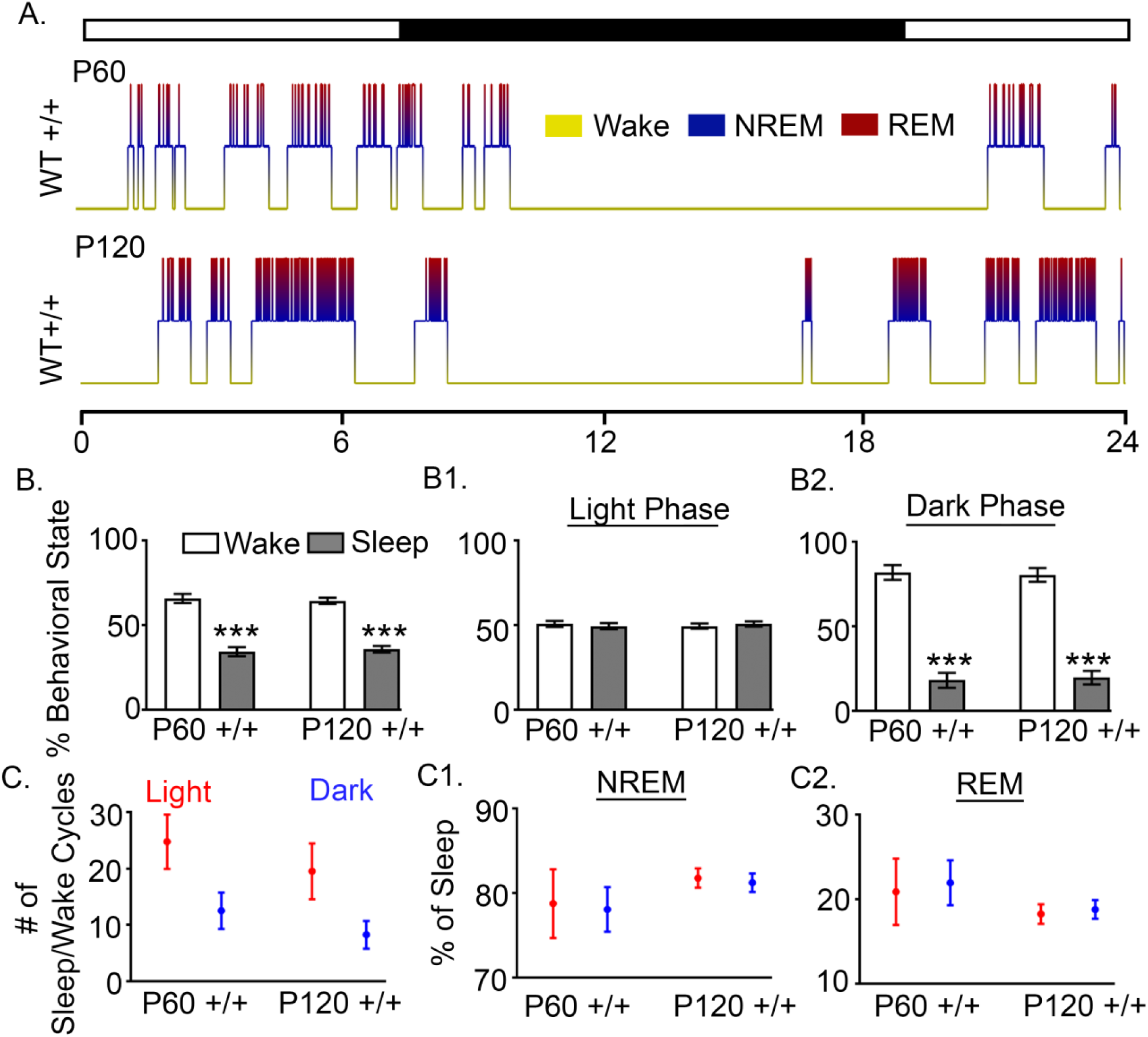
No differences in macro- or micro-sleep architecture of WT^+/+^ mice between the ages of P60 and P120. **(A)** 24h hypnograms of ultradian rhythms over the diurnal light/dark phase in WT^+/+^ mice at P60 (P60^+/+^) and P120 (P120^+/+^). **(B)** P60^+/+^ and P120^+/+^ mice spent significantly more time awake than asleep over the 24h period (P60^+/+^ Wake vs. P60^+/+^ Sleep: 2-way ANOVA, F_1,12_=162.1, P<0.0001, p<0.0001; P120^+/+^ Wake vs. P120^+/+^ Sleep: 2-way ANOVA, F_1,12_=162.1, P<0.0001, p<0.0001). **(B1)** P60^+/+^ and P120^+/+^ mice both did not have any significant differences between wake and sleep durations during the light phase**. (B2)** P60^+/+^ and P120^+/+^ mice spent significantly more time awake than asleep during the dark phase (P60^+/+^ Wake vs. P60^+/+^ Sleep: 2-way ANOVA, F_1,12_=216.9, P<0.0001, p<0.0001; P120^+/+^ Wake vs. P120^+/+^ Sleep: 2-way ANOVA, F_1,12_=216.9, P<0.0001, p<0.0001). **(C)** P60^+/+^ and P120^+/+^ mice did not have significant differences in the number of sleep/wake cycles during light vs. dark phase. **(C1 & C2)** The micro-architecture for the percent of both NREM and REM cycles during sleep showed an 80-20% split respectively, similar to WT^+/+^ mice, was and not significantly different betweenP60^+/+^ and P120^+/+^ mice. (p<0.05 *, p<0.01 **; P120 p<0.001 ***, post-hoc Bonferroni).

**Supplemental Fig. 4.**
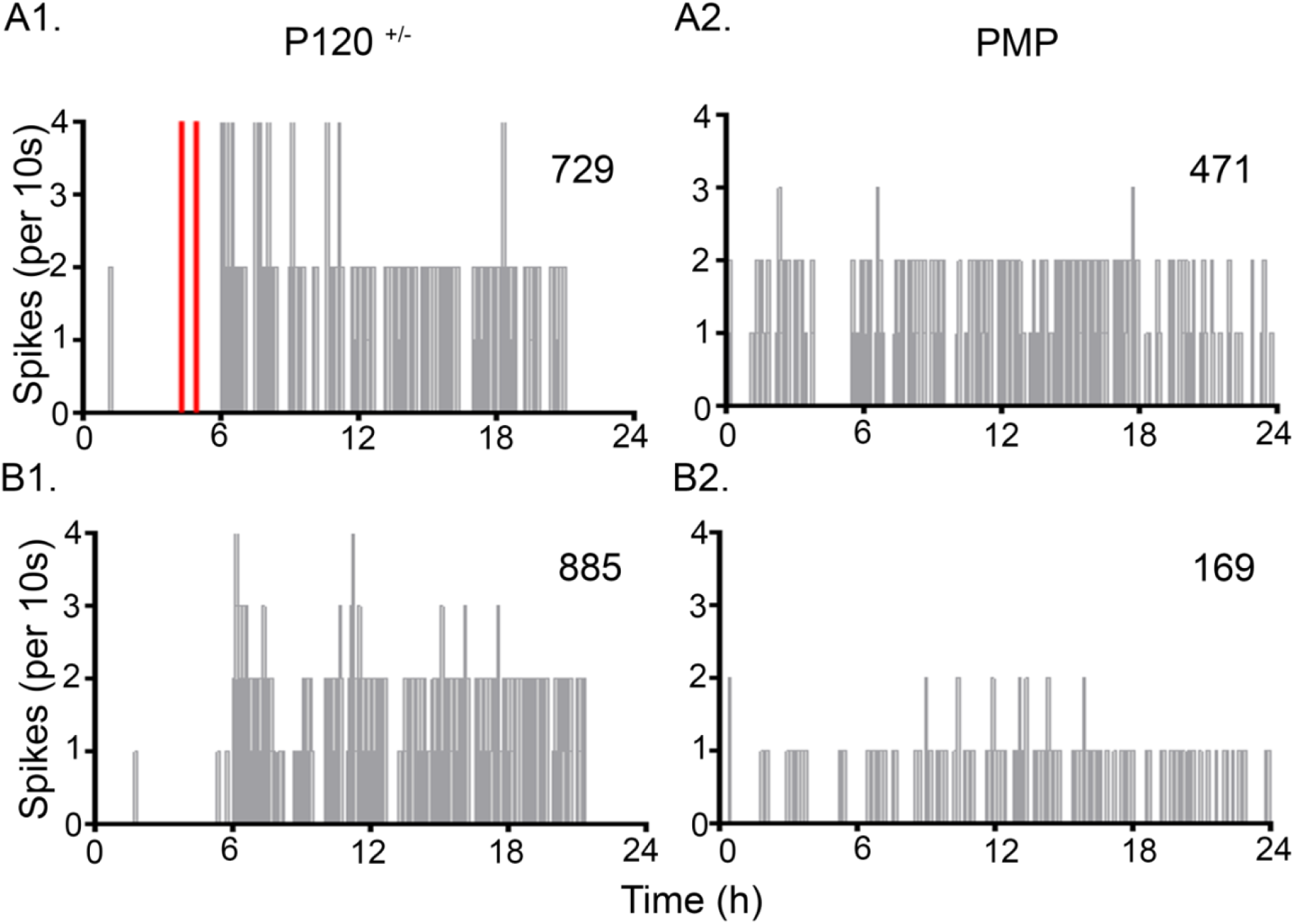
IISs frequency distributions before and after PMP administration. **(A1 & B1)** Representative frequency distributions of IIS (grey lines) and seizures (red lines) over 24h of EEG recording in two P120^+/-^ mice. **(A2 & B2)** Two doses of PMP (2mg/kg I.P.) administered during the EEG recording on P121 prevented seizure occurrence in *Syngap^+/-^* mice. PMP also reduced IIS rates (counts stated in each panel) which did not reach significance with the acute dosing protocol of PMP used in this study **(A2 & B2)**.

**Supplemental Fig. 5.**
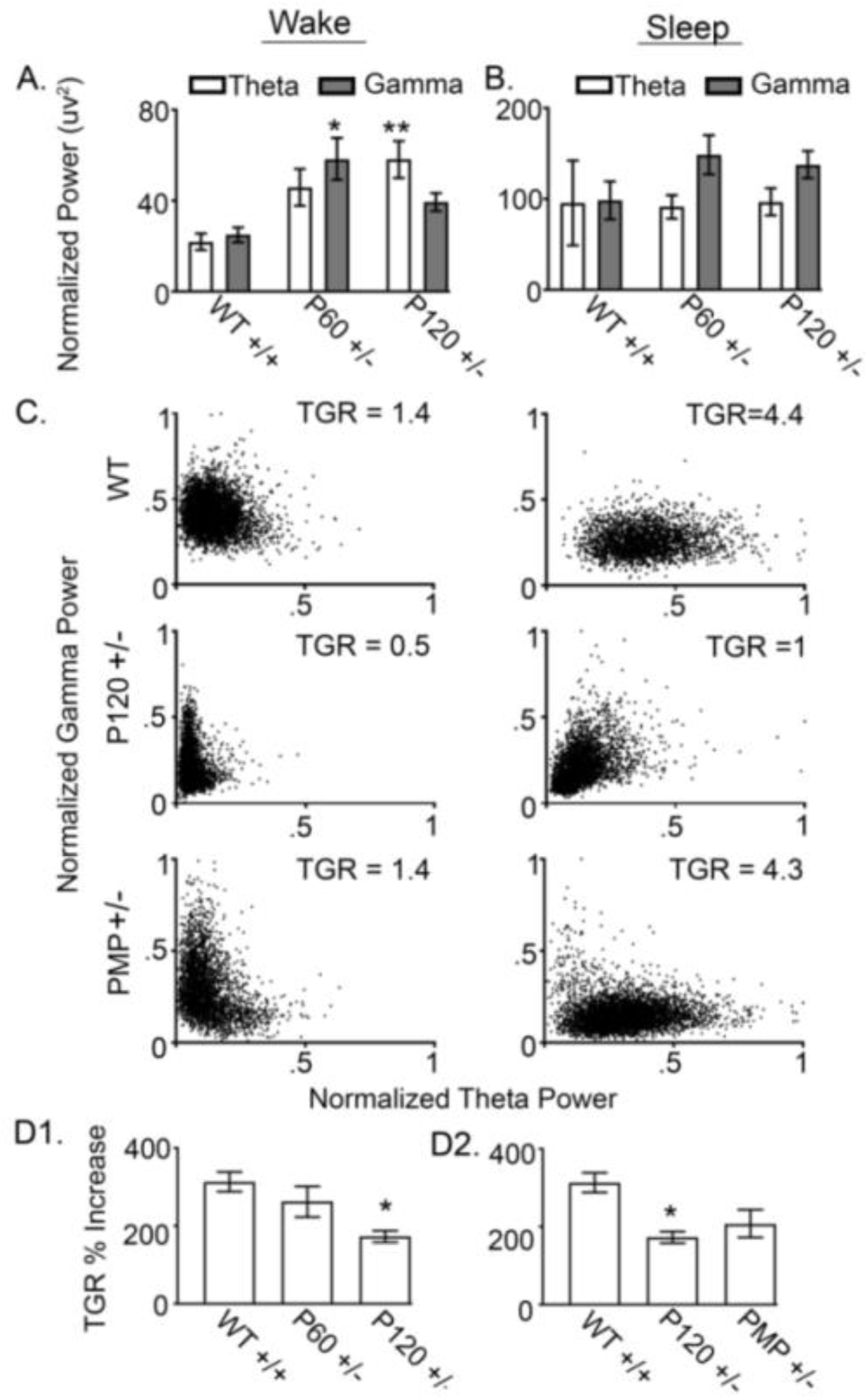
Dissociation of theta-gamma interaction during wake vs. sleep states. **(A)** 24h spectral power analyses of theta and gamma power during wake and sleep. Theta and gamma power during wake and sleep were normalized to their respective durations for each mouse. Theta power increased significantly during wake in P120^+/-^ mice (WT^+/+^ Wake Theta vs. P120^+/-^ Wake Theta: 1-way ANOVA, F_2,38_=11.64, P=0.0001, p=0.0028). Gamma power during wake was significantly increased in P60^+/-^ mice (WT^+/+^ Wake Gamma vs. P60^+/-^ Wake Gamma: 1-way ANOVA, F_2,38_=11.64, P=0.0001, p=0.021); this significance was lost in P120^+/-^ mice when compared to WT^+/+^ mice. **(B)** During sleep, there were no significant differences in either theta or gamma power compared to WT ^+/+^ mice. However, during sleep a general increase in gamma power was apparent in P60^+/-^ and P120^+/-^ mice. **(C)** Representative 24h theta-gamma ratio plots during wake and sleep in a WT^+/+^, P120^+/-^, and PMP-administered (2mg/kg I.P. inj; B.D.) mouse. WT^+/+^ mice had low theta power with high gamma power during wake states, resulting in a low theta-gamma ratio (TGR= 1.4). During sleep, WT^+/+^ mice had high theta power with low gamma power, resulting in a high TGR during sleep (TGR=4.4). P120^+/-^ mice demonstrated high gamma during both wake and sleep, resulting in similar TGRs during both wake and sleep states. **(D1)** The change in TGR from wake to sleep states was significantly reduced in P120^+/-^ mice (WT^+/+^ TGR % vs. P120^+/-^ TGR%:1-way ANOVA, F_2,16_=6.456, P=0.0088, p=0.0076). (**D2**) *Syngap1^+/-^* mice treated with PMP were not significantly different than WT^+/+^ mice. (p<0.05 *, p<0.01 **; P120 p<0.001 ***, post-hoc Bonferroni).

**Supplemental Fig. 6.**
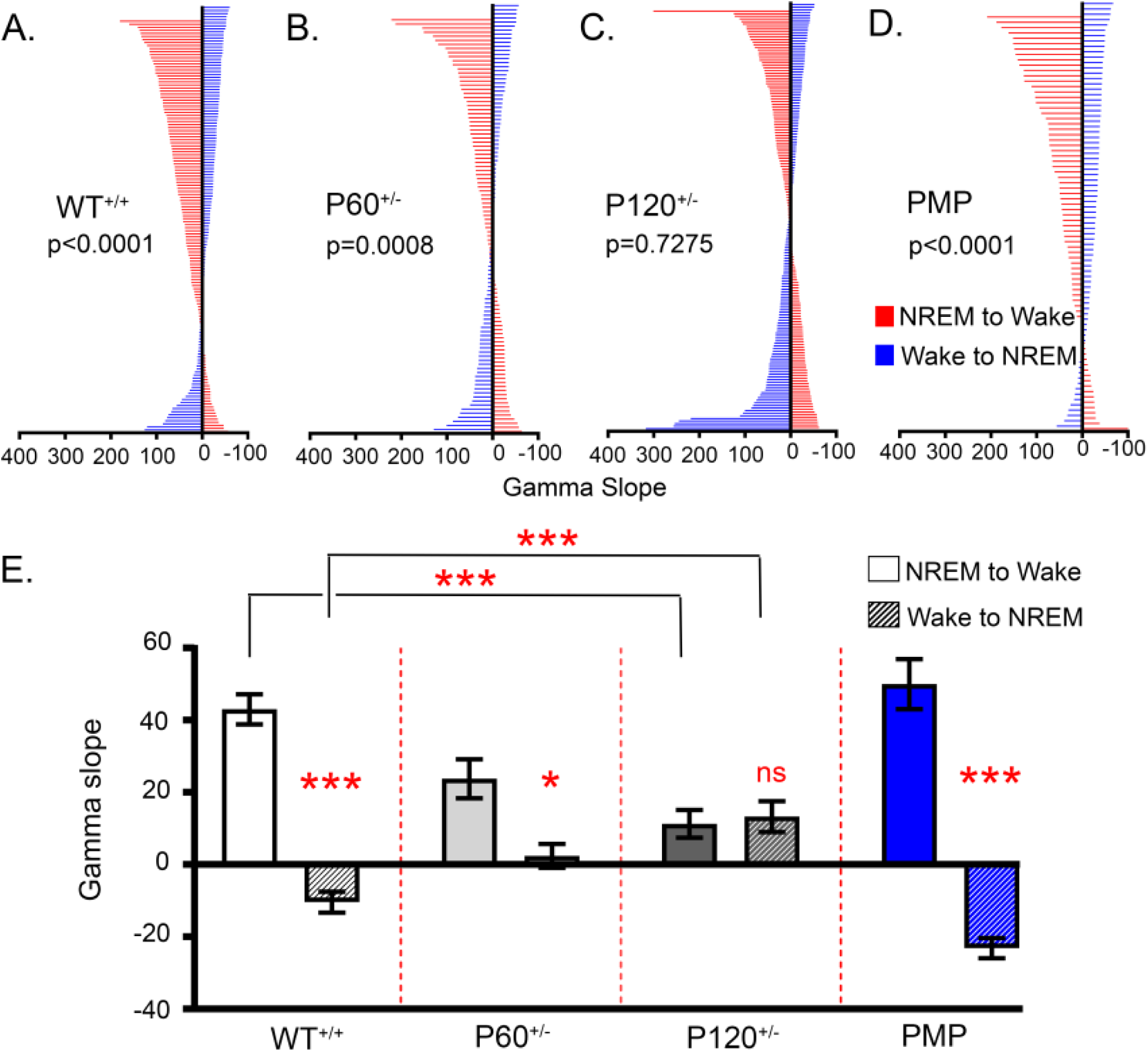
Frequency distribution of gamma slopes during Wake-NREM and NREM-Wake transitions. **(A)** The majority of gamma slopes during NREM-Wake transitions (red lines) were positive in WT^+/+^ mice, as gamma increased from NREM to wake. In contrast, the majority of gamma slopes during Wake-NREM transitions (blue lines) were negative, as gamma decreased from wake to NREM. In WT^+/+^, the slope of gamma during Wake-NREM and NREM-Wake transitions were state-dependent. **(B)** In P60^+/-^ mice, the differences between state-dependent gamma slopes were state-dependent. **(C)** In contrast, gamma slopes were not state-dependent in P120^+/-^ mice. **(D)** Administration of PMP restored state-dependent gamma slopes as the gamma slopes were positive during NREM-Wake transitions and negative during Wake-NREM transitions. **(E)** In WT^+/+^ mice mean gamma slopes for NREM-Wake and Wake-NREM transitions were significantly different (WT^+/+^ NREM to Wake vs. WT^+/+^ Wake to NREM: 2-way ANOVA, F_3,963_=28.24, P<0.0001, p<0.0001). Further, both NREM-Wake and Wake-NREM gamma slopes in WT^+/+^ mice were significantly different than those in P120^+/-^ mice (WT^+/+^ Wake vs. P120^+/-^ Wake: 2-way ANOVA, F_3,963_=28.24, P<0.0001, p<0.0001; WT^+/+^ NREM vs. P120^+/-^ NREM, p<0.0001). Gamma slopes of NREM-Wake and Wake-NREM transitions in P60^+/-^ mice remained significantly different (P60^+/-^ Wake vs. P60^+/-^ NREM: 2-way ANOVA, F_3,963_=28.24, P<0.0001, p=0.0313). In P120^+/-^ mice the gamma slope differences between NREM-Wake and Wake-NREM transitions were completely lost. Administration of PMP restored the significant differences between Wake-NREM and NREM-Wake transitions to WT^+/+^ levels (PMP^+/-^ Wake vs. PMP^+/-^ NREM: 2-way ANOVA, F_3,963_=28.24, P<0.0001, p<0.0001, post-hoc Bonferroni).

**Supplemental Fig. 7.**
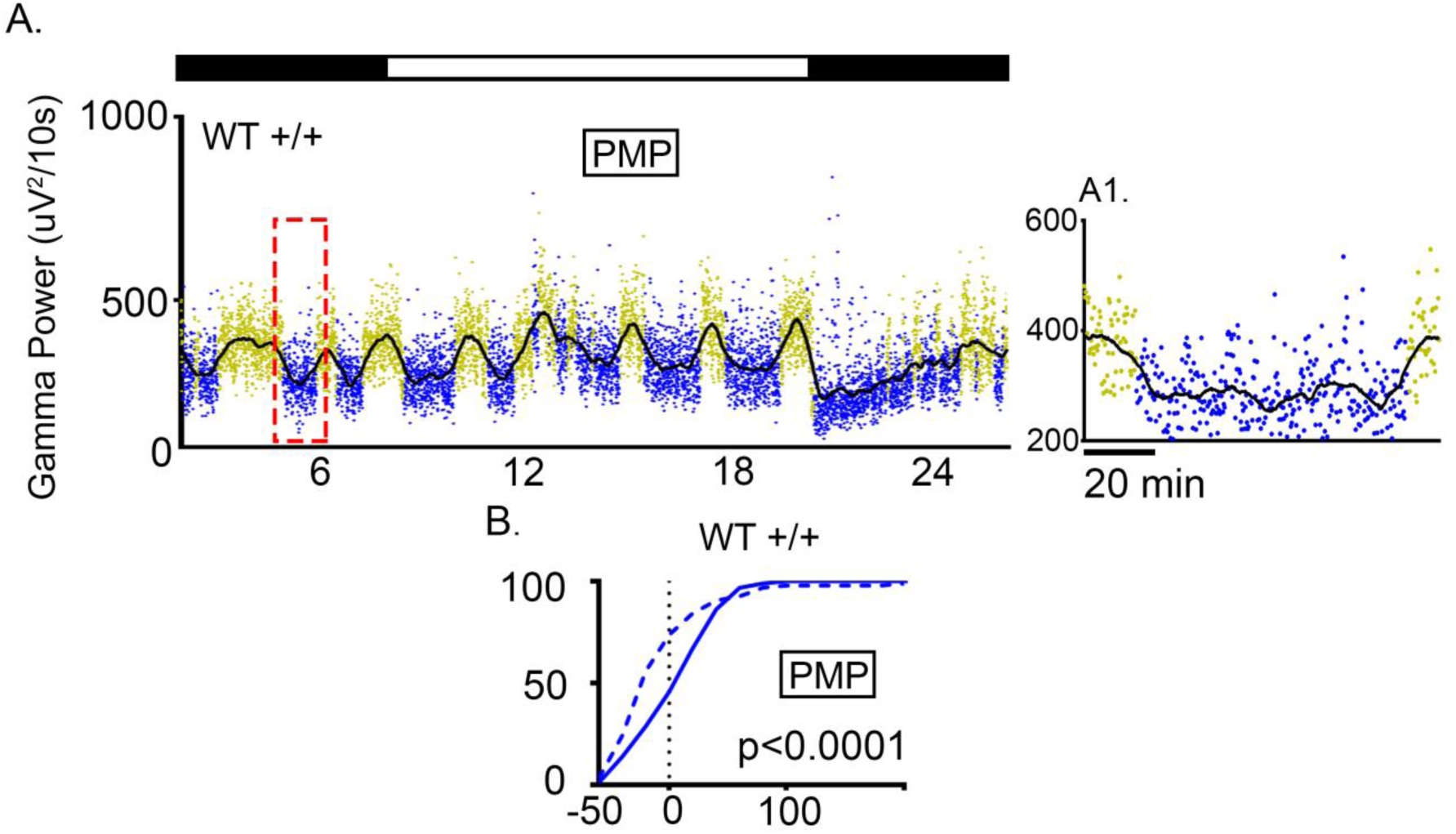
PMP treatment did not significantly alter cortical gamma spectral power transitions between wake and sleep at P120. **(A)** 24h WT^+/+^ mice treated with PMP have high gamma during wake and low gamma during NREM, similar to untreated WT^+/+^ mice (Fig. 7). **(A1)** Expanded time scales shows gradual fall of gamma during Wake-NREM transitions and gradual rise of gamma during NREM-Wake transitions**. (B)** Cumulative frequency plots of positive NREM to Wake slope (solid blue line) and negative Wake to NREM slope (dashed line), these slopes were significantly different (*t-test*, t_59_=10.5, p=0.0001). (p<0.05 *, p<0.01 **; P120 p<0.001 ***).

**Supplemental Fig. 8.**
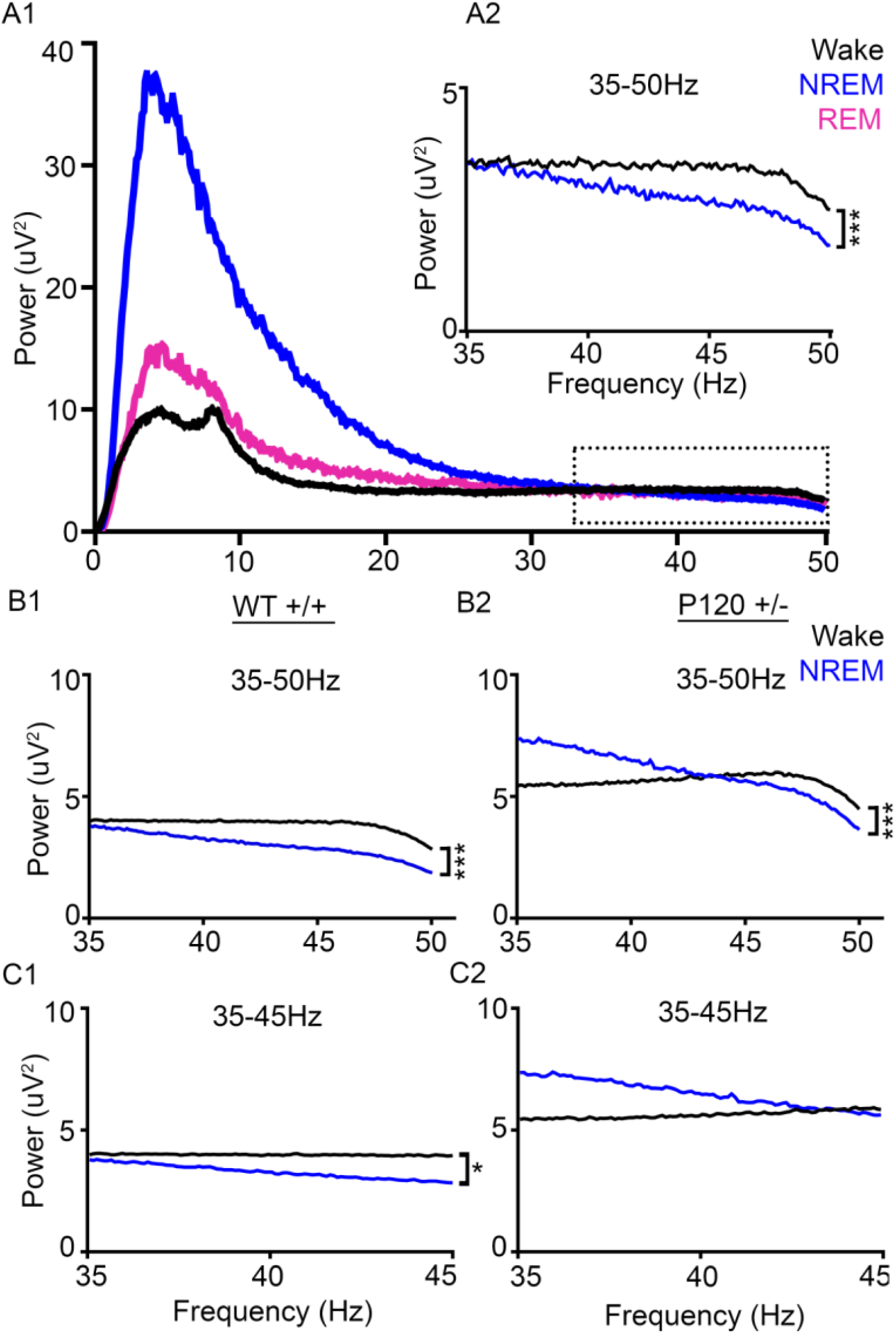
Gamma frequency analysis during wake and NREM. **(A1)** Respective frequency analysis (0.5-50Hz) of a WT^+/+^ mouse during wake, NREM, and REM. **(A2)** The gamma power (35-50Hz) of the respective WT^+/+^ mouse was significantly higher during wake than NREM (Wake vs. NREM: *t-test*, t_14.02_=300, P<0.0001). **(B1)** WT^+/+^ mice had significantly higher gamma power (35-50Hz) during wake than NREM (WT^+/+^ Wake vs. WT^+/+^ NREM: *t-test*, t_19.65_=300, P<0.0001). **(B2)** P120^+/-^ mice had significantly different gamma power from 35-50Hz during NREM than wake (P120^+/-^ Wake vs. P120^+/-^ NREM: *t-test*, t_4.25_=300, P<0.0001). **(C1)** Lower frequency gamma from 35-45Hz in wake and NREM was significantly different in WT^+/+^ mice (WT^+/+^ 35-50Hz Wake vs. WT^+/+^ 35-50Hz NREM: *t-test*, t_24.84_=200, P<0.0001). **(C2)** In P120^+/-^ mice, lower gamma frequency (35-45Hz) remained high during NREM. P120^+/-^ mice did not have any significant differences in gamma power (35-45Hz) between wake and NREM (P120^+/-^ 35-50Hz Wake vs. P120^+/-^ 35-50Hz NREM: *t-test*, t_15.31_=198, P<0.0001). (p<0.05 *, p<0.01 **; P120 p<0.001 ***).

**Supplemental Fig. 9.**
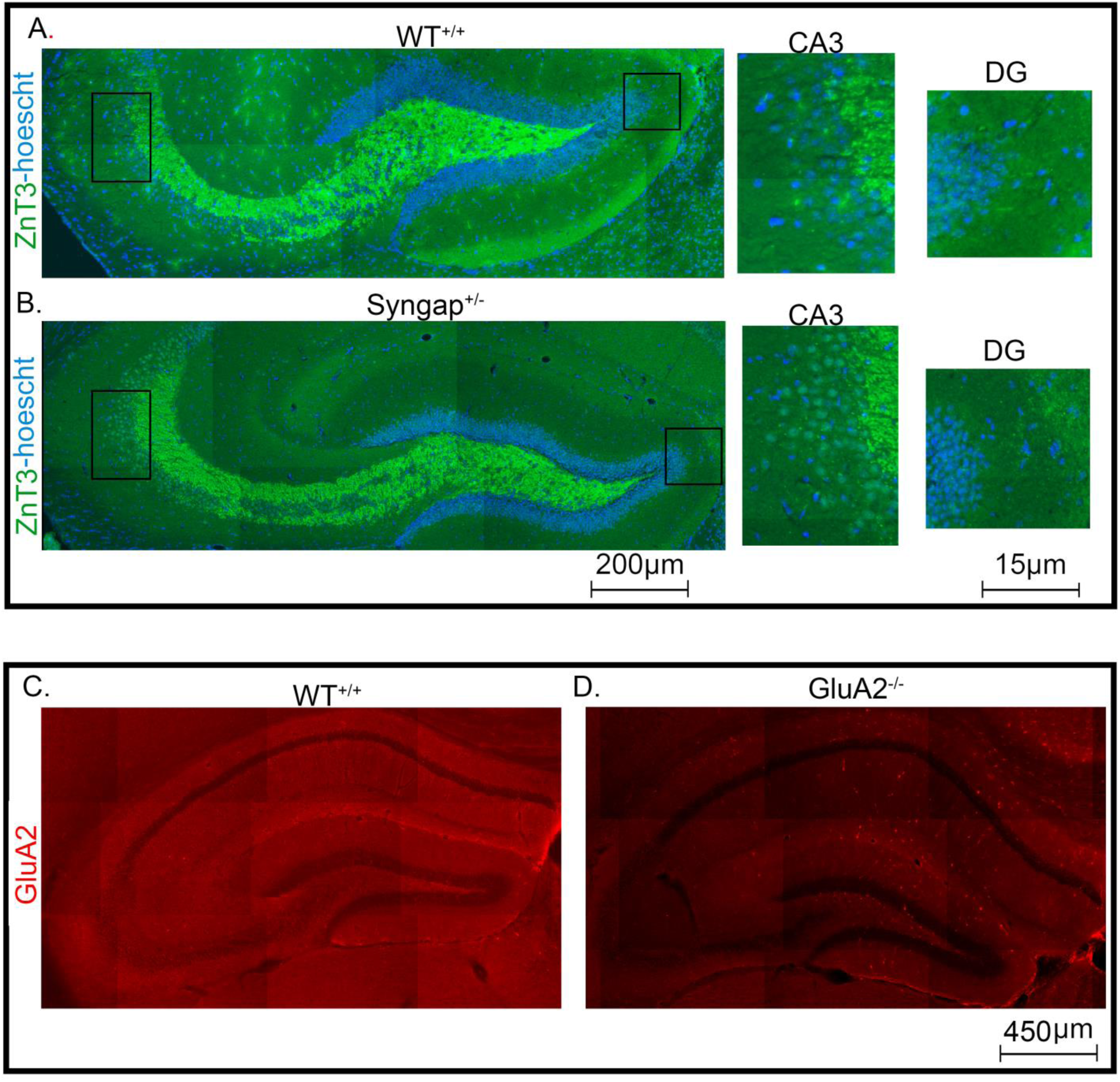
*Syngap1^+/-^* mice did not show evidence for mossy fiber sprouting and GluA2 antibody validation for IHC. **(A)** Representative 10X hippocampal section of a WT**^+/+^** mouse with ZnT3 staining (green) to label mossy fibers, an established marker of hippocampal epileptogenesis, with representative expansions of CA3 (left) and dentate gyrus (DG; right). **(B)** Representative 10X hippocampal section of a P120^+/-^ mouse demonstrated the absence of mossy fiber sprouting in both the DG and CA3. (C) Representative 10X hippocampal section of a WT^+/+^ mouse with GluA2 positive fluorescence and a GluA2^-/-^ mouse (Richard Huganir, Johns Hopkins University) lacking GluA2 fluorescence.

